# Telomere-to-telomere genome assembly of *Phaeodactylum tricornutum*

**DOI:** 10.1101/2021.05.04.442596

**Authors:** Daniel J. Giguere, Alexander T. Bahcheli, Samuel S. Slattery, Rushali R. Patel, Martin Flatley, Bogumil J. Karas, David R. Edgell, Gregory B. Gloor

## Abstract

*Phaeodactylum tricornutum* is a marine diatom with a growing genetic toolbox available and is being used in many synthetic biology applications. While most of the genome has been assembled, the currently available genome assembly is not a completed telomere-to-telomere assembly. Here, we used Oxford Nanopore long reads to build a telomere-to-telomere genome for *Phaeodactylum tricornutum*. We developed a graph-based approach to extract all unique telomeres, and used this information to manually correct assembly errors. In total, we found 25 nuclear chromosomes that comprise all previously assembled fragments, in addition to the chloroplast and mitochondrial genomes. We found that chromosome 19 has filtered long-read coverage and a quality estimate that suggests significantly less haplotype sequence variation than the other chromosomes. This work improves upon the previous genome assembly and provides new opportunities for genetic engineering of this species, including creating designer synthetic chromosomes.

## INTRODUCTION

*Phaeodactylum tricornutum* is a marine diatom that is described as a “diatom cell factory” (Butler et al., 2020) because it can be used to manufacture valuable commercial products. Recent genetic toolbox expansions, such as delivering episomes by bacterial conjugation (Karas et al., 2015) CRISPR-editing tools (Russo et al., 2018; Moosburner et al., 2020; Sharma et al., 2018; Stukenberg et al., 2018; Slattery et al., 2018; Serif et al., 2018), generation of auxotrophic strains (Zaslavskaia et al., 2000; Buck et al., 2018; Slattery et al., 2020), and the identification of highly active endogenous promoters (Erdene-Ochir et al., 2019) that are enabling rapid implementation of new product designs into commercial-scale production. The genome of *P. tricornutum* CCAP 1055/1 was sequenced in 2008, and resulted in a scaffold-level assembly predicting 33 chromosomes (NCBI assembly ASM15095v2) (Bowler et al., 2008). Chloroplast and mitochondrial genomes have also been published (Oudot-Le Secq et al., 2007; Oudot-Le Secq and Green, 2011), and have previously identified as targets for genetic engineering(Cochrane et al., 2020), as well as other chromosomes (Karas et al., 2013). Although the Bowler *et al*., assembly contains several telomere-to-telomere chromosomes, many scaffolds have only zero or one telomere, suggesting they are either incomplete or fragments of another chromosome. More recent work identifying centromeric sequences (Diner et al., 2017) in *P. tricornutum* has suggested that there may be less than 33 chromosomes. To generate a telomere-to-telomere assembly of *P. tricornutum* CCAP 1055/1, we used a hybrid approach with ultra-long reads from the Oxford Nanopore minION platform and highly accurate short reads from the Illumina NextSeq platform. We also introduce a novel graph-based approach to manually resolve telomere-related assembly errors. This approach identifies all unique telomere sequences and we demonstrate how it can be applied to manually correct assembly errors adjacent to chromosome ends. The full structural context of the *P. tricornutum* genome provides additional information for potential synthetic biology applications to manipulate the genome of this diatom cell factory.

## METHODS

### Growth

*Phaeodactylum tricornutum* (Culture Collection of Algae and Protozoa CCAP 1055/1) was grown in L1 medium without silica at 18^*°*^ C under cool white fluorescent lights (75 mE m^*−*2^ s^*−*1^) and a photoperiod of 16 h light:8 h dark as described previously (Slattery et al., 2018).

### DNA extraction

200 mL of culture (approximately 5 x 10^8^ cells) was spun at 3000 X g for 10 minutes at 4° C. The pellet was resuspended in 1 mL TE (pH 8.0) and added dropwise to a mortar (pre-cooled at -80° C) pre-filled with liquid nitrogen. The frozen droplets were ground into a fine powder with a mortar and pestle, being careful to keep the cells from thawing by adding more liquid nitrogen as necessary. The frozen powder was transferred to a 15 mL Falcon tube where 2 mL of lysis buffer was added (1.4 M NaCl, 200 mM Tris-HCl pH 8.0, 50 mM EDTA, 2% (w/v) CTAB, RNAse A (250 *µ*g/mL) and proteinase K (100 *µ*g/mL)). The solution was mixed very slowly by inversion, incubated for 30 minutes at 37° C (mixed very slowly halfway through incubation). Cellular debris was pelleted at 6000 X g for 5 minutes. Lysate was transferred to a new 15 mL Falcon tube. One volume of 25:24:1 phenol:chloroform:isoamyl alcohol was added, mixing slowly by inversion. The sample was centrifuged at 6000 X g for 5 minutes. The aqueous phase was transferred as slow as possible to a new Falcon tube. One volume of 24:1 chloroform:isoamyl alcohol was added, and mixed slowly with end-over-end inversion. The sample was centrifuged at 6000 X g for 5 minutes. Approximately 450 uL of the aqueous phase was transferred into new 1.5 mL Eppendorf tubes. To the Eppendorf tube, 1/10 volume of 3 M NaAc pH 5.2 and 2 volumes (final volume) of ice-cold 100% ethanol were added, mixing slowly by end-over-end inversion. The sample was centrifuged at 16 000 X g for 5 minutes, and washed twice with 500 uL 70% ethanol. Ethanol was decanted, and the pellet was dried for approximately 10 minutes by inverting on a paper towel. The pellet was resuspended in 100 uL 10 mM Tris-HCl pH 8.0, 0.1 mM EDTA pH 8.0. After resuspending overnight at 4° C, short DNA fragments were then selectively removed using the Short Read Eliminator (SRE) kit from Circulomics (Baltimore). DNA from the same extraction was used for sequencing on both the Oxford Nanopore minION and Illumina NextSeq 550 platform.

### Sequencing

All sequencing reads are publically availabe on the European Nucleotide Archive under project PR-JEB42700. All raw .fast5 files are available on under accession number ERR5858460.

An Oxford Nanopore minION flow cell R9.4.1 was used with the SQK-LSK109 Kit according to the manufacturer’s protocol version GDE 9063 v109 revK 14Aug2019, with one alteration: for DNA repair and end-prep, the reaction mixture was incubated for 15 minutes at 20° C and 15 minutes at 65° C. Basecalling was performed after the run with Guppy (Version 3.6). NanoPlot (De Coster et al., 2018) was used to generate Q-score versus length plots and summary statistics. The read N50 of the unfiltered reads was approximately 35 kb (Fig. S1). Nanopore reads are available under accession ERR5207170.

For Illumina sequencing, the Nextera XT kit was used to prepare 2×75 paired-end mid-output NextSeq 550 run at the London Regional Genomics Center (lrgc.ca). Reads were trimmed using Trimmomatic v0.36 (Bolger et al., 2014) in paired end mode with the following settings: AVGQUAL:30 CROP:75 SLIDINGWINDOW:4:25 MINLEN:50 TRAILING:15. SLIDINGWINDOW AND TRAILING were added to remove poor quality base calls. Only paired end reads were retained. Illumina reads are available under accession ERR5198869.

## Assembly

### Telomere identification

We first obtained sequences for the end of every linear chromosome. The sequence of the telomere repeats for *P. tricornutum* are known from the previous assembly (Bowler et al., 2008) to be repeats of AACCCT. All long reads larger than 50 kilobases with 3 or more consecutive telomeric repeats (or the reverse complement) were extracted by filtering using NanoFilt (De Coster et al., 2018) and by string matching using grep. All-versus-all mapping of the telomeric reads was performed using minimap2 (Li, 2016). Only overlapping reads with a minimum query coverage of 95 % were retained.

To determine the sequence of unique telomeres for each chromosome, a network graph was generated with iGraph (Csardi and InterJournal, 2006). Each read name was used as a vertex, and edges were generated between each overlapping read with more than 95% query coverage. Noise was filtered by removing any group of overlaps with less than 5X coverage. There were 95 vertices that had greater than 20X coverage; that is, there are 95 unique telomere sequence groups. Most groups had approximately 40X coverage (number of long reads per group), however, several outliers had with more than 60X coverage. These represent duplicated regions in the telomeres that are not unique (i.e., more than one haplotype or chromosome contains this sequence). We estimate that ninety-five unique telomere groups indicates that there are at least 24 or more chromosomes (95 chromosomes divided by 2 ends per chromosome divided by 2 haplotypes). There are at least 88 groups of telomere sequences that are unique and can be used to improve the assembly, with the remaining 7 possibly duplicated. The longest read of each telomere group, typically greater than 100 kb in length, was retained as a representative telomere sequence for correction. Example code for this is available in Code S7.

### Assembly

Several recent assembly algorithms and with multiple parameters were attempted, but we found overlap-layout consensus to provide the most contiguous assembly as a starting point. Sequencing reads longer than 75 kilobases were used for initial assembly with miniasm, (Li, 2016) using the parameters -s 30000 -m 10000 -c 5 -d 100000. From this initial assembly, unitigs were manually completed with the following approach:

1) Mapping of telomeric reads against the unitig. If no telomere was present on the unitig and a high query coverage alignment was found, the unitig was extended to the telomere sequence of the mapped telomere. 2) After telomere extension (or confirmation), reads longer than 50 kb were mapped to the unitig to confirm overlapping coverage over the entire chromosome. Coverage was evaluated using only reads larger than 50 kb and with higher than 60% query coverage, with an alignment score:length ratio less than 2 (similar to previous validation methods)(Giguere et al., 2020). A query coverage of only 60% was chosen to allow for potential haplotype divergence. 3) Telomere-to-telomere unitigs with overlapping ultra-long read coverage and no gaps were deemed validated and brought forward to improve base accuracy by read polishing.

The chloroplast and mitochondrial genomes were assembled using a reference based approach by first extracting all reads that aligned to the reference chloroplast and mitochondria with high query coverage. Reads were then *de-novo* assembled using miniasm.

### Polishing

Due the repetitive nature of the genome and the diploid nature of *P. tricornutum*, raw assemblies were polished using an iterative approach with racon (Vaser et al., 2017), medaka (Oxford Nanopore) and Pilon (Walker et al., 2014) as described in the Methods S6. Briefly, after each polishing iteration, we corrected errors that were introduced by the polishing algorithms as described in Methods S6, and modified the medaka polishing by filtering reads using a minimum of 50% query coverage. The assembly was first polished by nanopore reads only, followed by Illumina read polishing using Pilon. For the chloroplast and mitochondria, the subset of reads identified as either chloroplast or mitochondria were used for polishing.

### Methylation

5mC methylation sites were predicted using Megalodon v2.2.1 (Oxford Nanopore Technologies) using the model res dna r941 min modbases 5mC CpG v001.cfg from the Rerio repository (Oxford Nanopore Technologies) with Guppy 4.5.2. A default threshold of 0.75 was used as a minimum score for modified base aggregation (probability of modified/canonical base) to produce the final aggregated output. The percentage of reads methylated at the predicted are plotted in Fig. S2.

All data and code used to generate figures are available from DOI: 10.5281/zenodo.4731048.

## RESULTS

### Workflow

We developed a sample and library preparation protocol that provided exceptionally long reads. We observed a read N50 of 35 kilobases, with the longest reads just over 300 kb, following sequencing with the Oxford Nanopore minION platform. Of the 7.8 gigabases of raw sequence data, approximately 2.5 gigabases were from reads longer than 50 kilobases Figure S1). We found that chromosomes assembled using standard approaches were often mis-assembled around telomeres, or were fragmented and only contained 1 telomere. To correct each unitig, we leveraged the unique ultra-long telomere reads as described in the Methods S6 and in Figure 1. This approach was used to manually identify a tiling path for each chromosome until each chromosome was contiguous from telomere to telomere and validated by a tiling overlapping read path.

**Figure 1.**
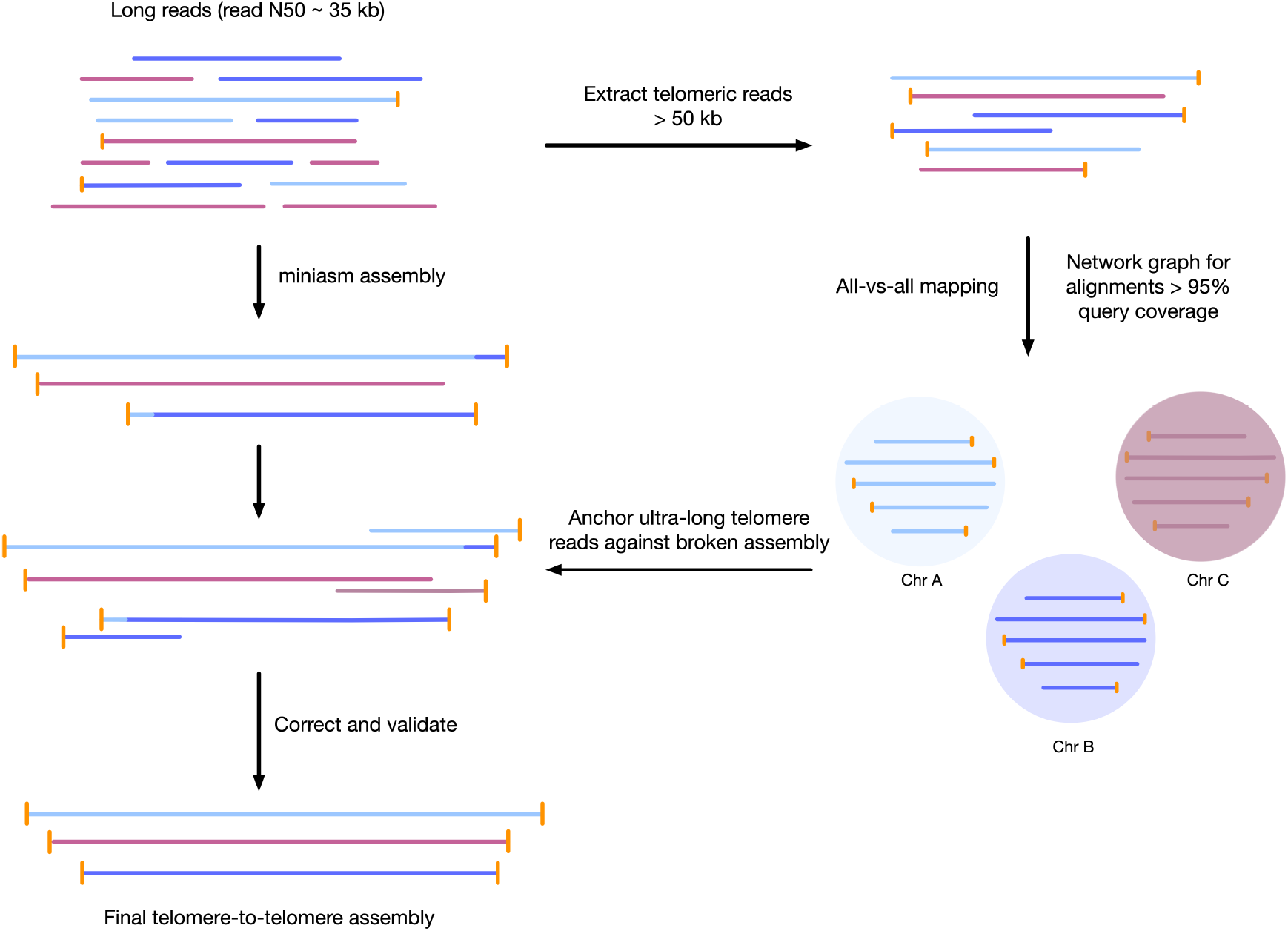
Workflow for telomere-to-telomere genome assembly. Telomere-containing nanopore reads larger than 50 kb are extracted, and are mapped in all-vs-all mode using minimap2. The resulting alignments are filtered by 95 % query coverage, and a network graph is created using iGraph using read names as vertices, and alignments between reads as edges. Each resulting cluster represents one end of a chromosome. The initial reads are used for overlap-consensus assembly using miniasm. On a chromosome-by-chromosome basis, ultra-long read coverage is plotted. If an assembled chromosome is missing a telomere or has an assembly error revealed by a lack of overlapping read coverage, the longest read from each telomere cluster is mapped against the chromosome, and the resulting telomere is used to manually correct the assembly and extend to the telomere using an overlap-layout consensus approach.

### Tiling path of overlapping reads verify contiguity

To ensure our genome assembly is contiguous, we generated multiple independent minimum tiling paths of overlapping long reads (Data S5, Fig. S2). Reads larger than 50 kb were mapped against the assembly using minimap2. To ensure no incorrect alignments were retained, any reads with less than 90% of the read aligned to the assembly were removed. From this subset, 5 independent minimum tiling paths that required at least 10 kb of overlap between each read was generated. All chromosomes have multiple independent (i.e., no common reads) tiling path of reads with a minimum overlap of 10 kb in the final validated assembly (5 independent paths shown in .paf format in Data S5), indicating that all chromosomes are contiguous. Chromosomes were manually corrected to meet this standard if necessary.

In addition to overlapping reads, Fig. S2 also shows the GC content for each chromosome. A previous study has proposed that centromeres could be identified by low GC content (Diner et al., 2017). The 100 base window with the minimum GC content are shown in Fig. S2 and are highlighted between red dotted lines. Putative centromere sequences characterized by the lowest GC content window as previously described Diner et al. (2017) are highlighted between in red.

### Telomere-to-telomere assembly comprises previous scaffolds

We ultimately obtained 25 telomere-to-telomere chromosome assemblies that recruit 98% of long reads, and these chromosomes comprised all previously proposed chromosomes from Bowler *et al*., (2008), as well as circularized chloroplast and mitochondrial genomes. The median coverage for unfiltered long reads across the nuclear genome was 202X, while median coverage for the chloroplast and mitochondrion were approximately 6201X and 528X, respectively. This was calculated in 1000 base windows using mosdepth (Pedersen and Quinlan, 2018).

A key feature of this updated assembly is the consistency with previous sequencing efforts (Bowler et al., 2008). Previously, 25 centromere sequences were identified (Diner et al., 2017), suggesting that there were fewer than the proposed 33 chromosomes. This agrees with our conclusion of 25 nuclear chromosomes. In addition, we independently resolved the relative location for all of the previously proposed partial chromosomes without internal inconsistencies in Figure 2 (i.e., scaffolds with only 1 telomere were resolved into a telomere-to-telomere chromosome).

**Figure 2.**
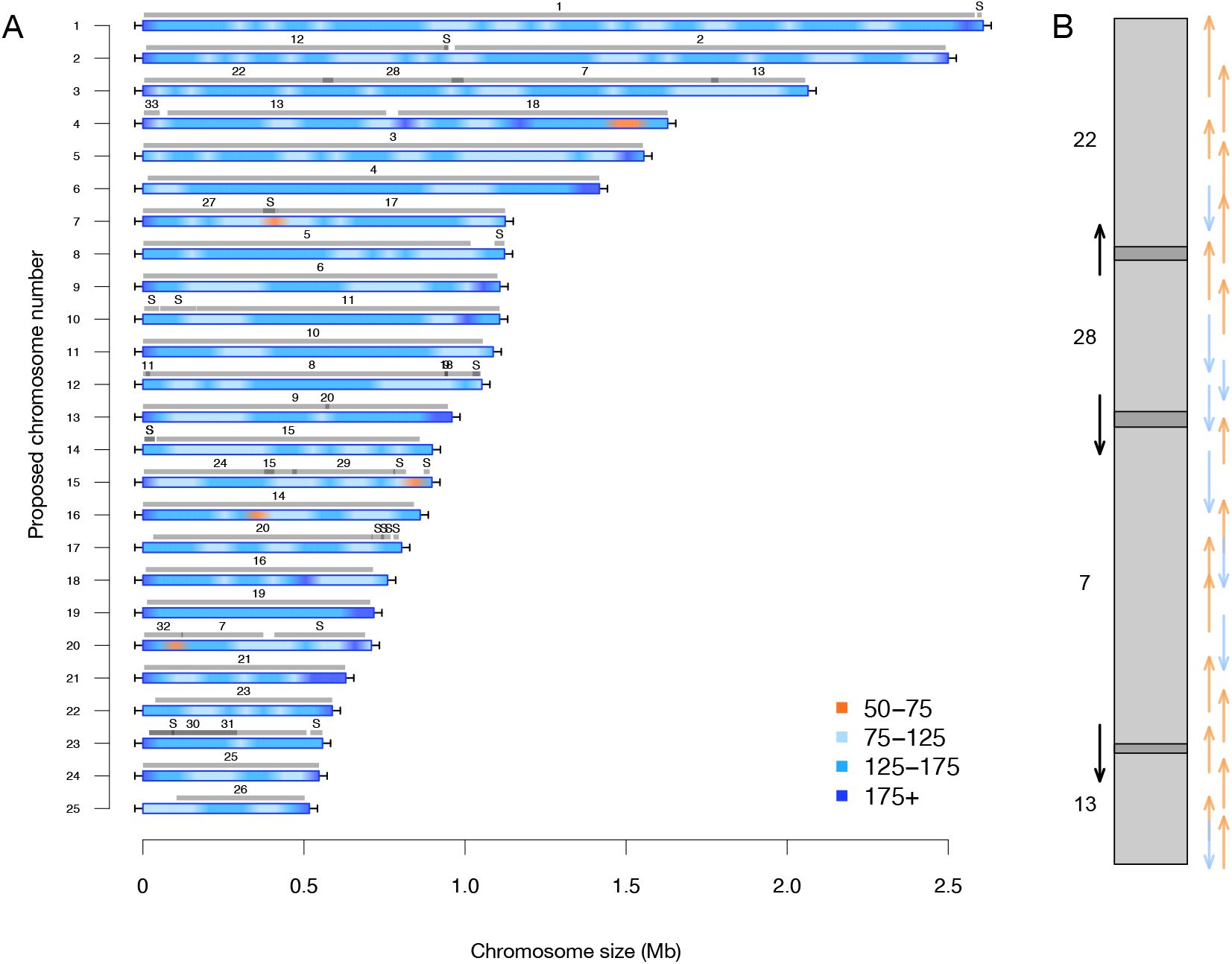
A) Filtered long-read coverage and comparison to previous assembly. Reads longer than 20 kb were mapped against the assembly, filtered (minimum 20000 base alignment and 50 % query coverage), and genome coverage was calculated in 50 kb windows using mosdepth. Newly proposed chromosomes names are left (by length). Scaffolds from the previous genome assembly (ASM15095v2) are overlayed as grey bars, aligned using minimap2 in asm5 mode and filtered to retain minimum 10 kb alignments. Numbers on top of gray bars indicate which previous scaffold number, with S representing small unassembled scaffolds. Horizontal “T” bars on each end indicate telomere-repeat presence. B) Visualization of proposed chromosome 3 with alignments to previous chromosomes. Dark gray regions indicate overlap. Coloured arrows on the right indicate minimum overlapping read path (orange = negative strand, blue = positive strand, black arrows on left show ultra-long reads that completely span regions where previous assembly could not assemble through.

### Assembly quality

To assess the quality of the assembly, we used Merqury (Rhie et al., 2020) to estimate the base-level accuracy and completeness by k-mer frequency, shown in Data S3. We found that the estimated quality value (estimated log-scaled probability of error for the consensus base calls by Merqury) ranged from 27 - 53, depending on the chromosome. The mean quality value for nuclear chromosomes was 28.86, with chromosome 19 as an outlier at 43. The QV for all nuclear genomes except for 19 are likely lower because the chromosomes were polished using reads that are heterozygous - this can likely be improved in the future by binning the reads into haploypes before polishing. The chloroplast and mitochondrial genomes have a quality value of 53 and 42, respectively. Importantly, the k-mer completeness estimate of 80% suggests that many k-mers in the Illumina reads are not represented in this genome assembly, implying there is still additional genomic sequence missing from this assembly. This was also the case when using the Bowler assembly as input for Merqury. This is expected because this assembly is not yet haplotype-resolved.

We also show that nearly all the reads are recruited in Table 1. When reads are filtered by query coverage, the number of reads recruited drops to 74%, indicating there is still sequence information to be determined by resolving the haplotypes.

**Table 1.** Long reads were filtered to remove all reads smaller than 1000 bases and below a Q-score of 8 using NanoFilt. Lambda spike-in reads were then removed used NanoLyse, and the total number of reads was calculated using NanoPlot. The number of reads recruited was calculated by aligning the reads against the assembly using minimap2, and unique read ideas were counted. The number of filtered reads was counted after removing reads with less than 90 percent query coverage.

| Unique reads recruited | Unique filtered reads recruited |
| --- | --- |
| 98.12% | 74.00% |

### Filtered long-read coverage for Chromosome 19 is inconsistent with diploid state

We observed that chromosome 19 has remarkably consistent (i.e., no drops in coverage) filtered long-read coverage relative to the other linear chromosomes (Figure 2, Fig. S2). *Phaeodactylum tricornutum* is a diploid organism but has not been observed to undergo meiosis. Therefore, we expected to observe 2 unique haplotypes per chromosome. These 2 haplotypes can be inferred by drops in filtered-long read coverage to half the total coverage, which indicates that the chromosome is a combination of both haplotypes (Fig. S2). This variation in coverage is observed for all nuclear chromosomes with the striking exception of chromosome 19. Chromosome 19 also has duplicated coverage near the telomeres. These observations suggest that there are not two haplotypes for chromosome 19, suggesting a different recent history for this chromosome in this strain.

### 5mC methylation and transposable elements

The Extensive de-novo TE Annotator (EDTA) pipeline (Ou et al., 2019) was used to predict transposable elements in the genome. We found that the majority of transposable elements are long terminal repeat (LTR) retrotransposons (3.64% of the genome was found to be Copia-type, 5.65% were unknown - Data S4) while terminal inverted repeats were only 1.17% of the genome, and helitrons were 0.54% of the genome. Each LTR region is represented as shaded blue regions in Fig. S2 in blue, and density plots of the end locations are shown in the top quadrant. Chromosome 19 contained the fewest transposable elements at 50. The locations and density of LTR-retrotransposons are plotted in Figure 3 for proposed Chromosome 3 and Fig. S2 for all other chromosomes.

**Figure 3.**
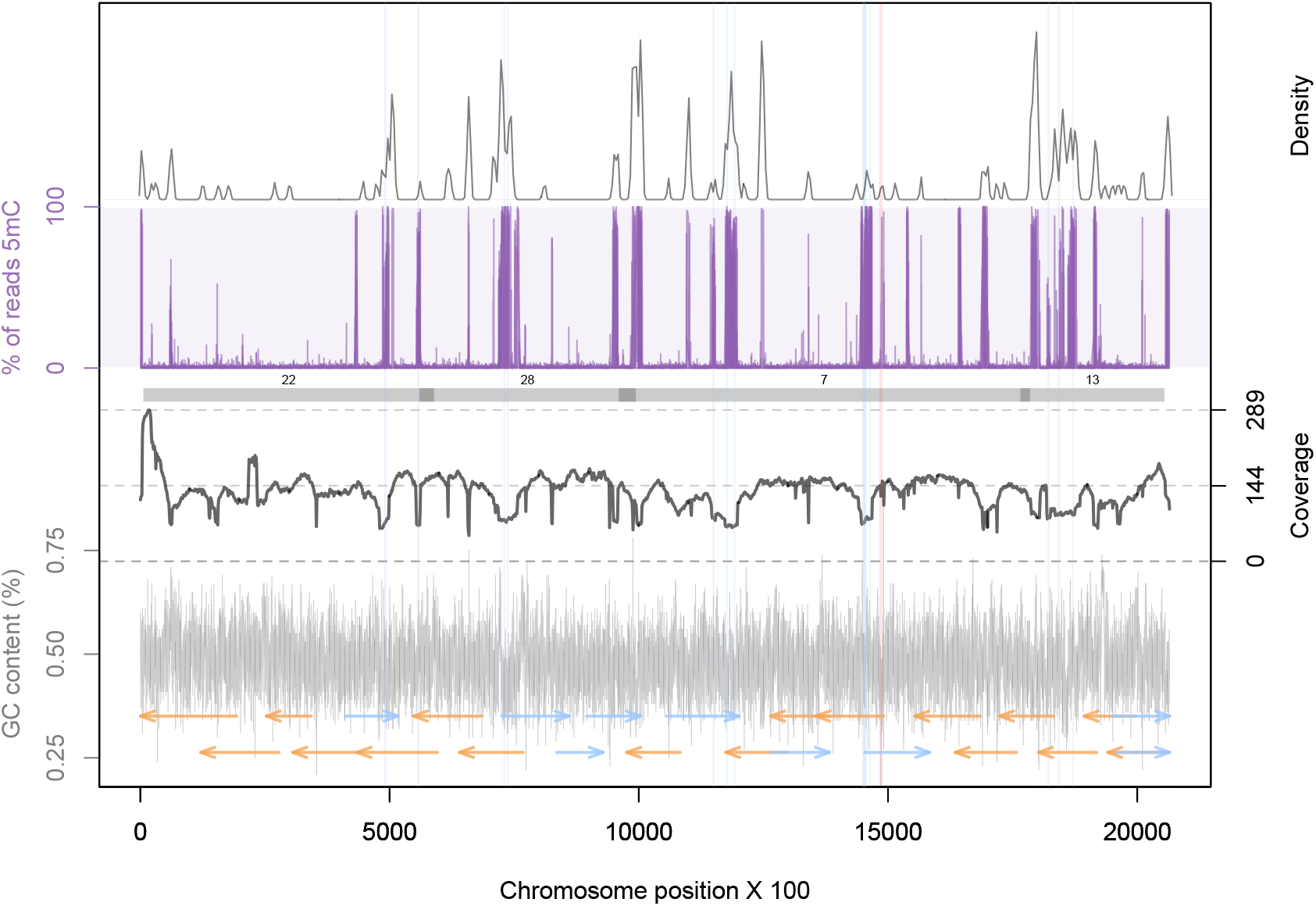
Genomic summary for proposed Chromosome 3. GC content was calculated in 100 base windows, and plotted in the bottom quadrant. An overlapping read tiling path, with a minimum overlap of 30 kb, is shown with orange indicating reads mapping to the negative strand and blue indicating reads mapping to the positive strand. Filtered long-read coverage is plotted in the second quadrant (minimum 20 kb length and 70% query coverage). Number scaffolds from the previous assembly are overlayed in gray bars, with dark grey representing overlapping regions. The third quadrant shows the proportion of reads that were called as methylated. The top quadrant shows the density of LTR-retrotransposon end points as predicted by the EDTA pipeline. Light blue box (full plot) shows the regions that are annotated at LTR-retrotransposons. Light red box indicates the 100 base window with the minimum GC content.

Previous studies have found that some tranposable elements were hypermethylated (Veluchamy et al., 2013). Using chromosome scale nanopore methylation basecalling, we found a strong signal between many predicted LTR retrotransposons and methylation status (Figure 3, Fig. S2).

We examined the association between LTR transposon dense regions and the locations of low coverage regions and regions where the previous assembly failed to generate overlapping regions. We observed that scaffolds with overlapping regions (Fig. S2) generally were not assembled into full chromosomes because of ambiguity in the placement of the LTR-rich regions at the ends of the scaffolds. These are now resolved by the long-read assembly identified here. Additionally, many of the low-coverage regions of our assembly overlap with the locations of the LTR-dense regions and this would be consistent with chromosomal rearrangements being more likely in these regions. Further investigation at these regions is required.

## DISCUSSION

Here, we developed a graph-based approach to locate the unique telomere ends of all *P. tricornutum* chromosomes, and applied this information to generate an telomere-to-telomere assembly. The new assembly incorporates all the previous chromosome fragments from Bowler (2008), and also includes circularized organelle genomes.

The chromosomes show marked variations in genome coverage in a two-fold range. This suggests that there are large regions of the chromosomes that have substantial haplotype differences. Strikingly, one chromosome has completely consistent coverage between the telomeres. While this needs to be further investigated, we speculate that this chromosome in this strain may have undergone a recent sequence homogenization event. Previous work has also found that chromosome 19 appears homozygous in the wild type strain (Russo et al., 2018). It has previously been speculated that *Phaeodactylum tricornutum* may be capable of sexual reproduction (Mao et al., 2020; Patil et al., 2015), but there has yet to be conclusive evidence.

We also demonstrate that nanopore sequencing can identify methylated regions associated with transposable elements (Fig. S2). While this strain of algae was not grown under stress conditions, we demonstrate that this new technology can be applied to to characterize the 5mC methylome of *P. tricornutum*, and can be applied to future differential methylation experiments.

Chromosome 19 has a high quality value of 43, while the other nuclear chromosomes have lower quality values around 28, respresenting an expected drop in per-base quality due to polishing the assembly with a heterozygous read set. Taken together with the consistent filtered-long read coverage, these data suggest that there are not highly divergent haplotypes of chromosome 19. In addition, the higher quality values for the organelle genomes and chromosome 19 indicate that this dataset is sufficient to generate highly accurate (quality value greater than 40) assemblies when haplotypes can be resolved. Furthermore, the k-mer completeness estimate suggests there is additional unknown genomic information (up to 20% of k-mers in Illumina reads), and we believe the remaining sequence can be identified by generating a haplotype-resolved genome assembly for *P. tricornutum*. We have deposited all raw short and long-read data publicly for use by the community as Project PRJEB42700 on the European Nucleotide Archive.

While this work improves on the previous genome assembly, we believe there is yet further room to improve our understanding of the *P. tricornutum* genome. First, we believe that haplotype-phasing can be completed with ultra-long read data. Furthermore, there modified bases can be further explored in relation to gene expression. We have therefore posted the raw files and encourage the community to investigate other genomic features with this data. We also recognize that the per-base accuracy of this assembly may be improved by binning the reads used for polishing by haplotype - as evidence by the high quality value estimate for chromosome 19 and the organeller genomes. The per-base quality of this genome assembly will likely benefit from resolving the haplotypes and re-analysis as improvements to basecalling and polishing algorithms become available in the future. This telomere-to-telomere genome assembly will make it possible to start designing and creating synthetic chromosomes in *Phaeodactylum tricornutum*.

## FUNDING

DJG Mitacs number IT8360, Ontario Graduate Scholarship

SS: Mitacs number IT8360

GBG: Natural Sciences and Engineering Research Council of Canada (NSERC), grant number: RGPIN-03878-2015

BK: Natural Sciences and Engineering Research Council of Canada (NSERC), grant number: RGPIN-2018-06172

DE: Natural Sciences and Engineering Research Council of Canada (NSERC) Discovery Grant RPGIN-2015-04800

