## Supplemental Figure 2 for "Telomere-to-telomere genome assembly of *Phaeodactylum tricornutum*"

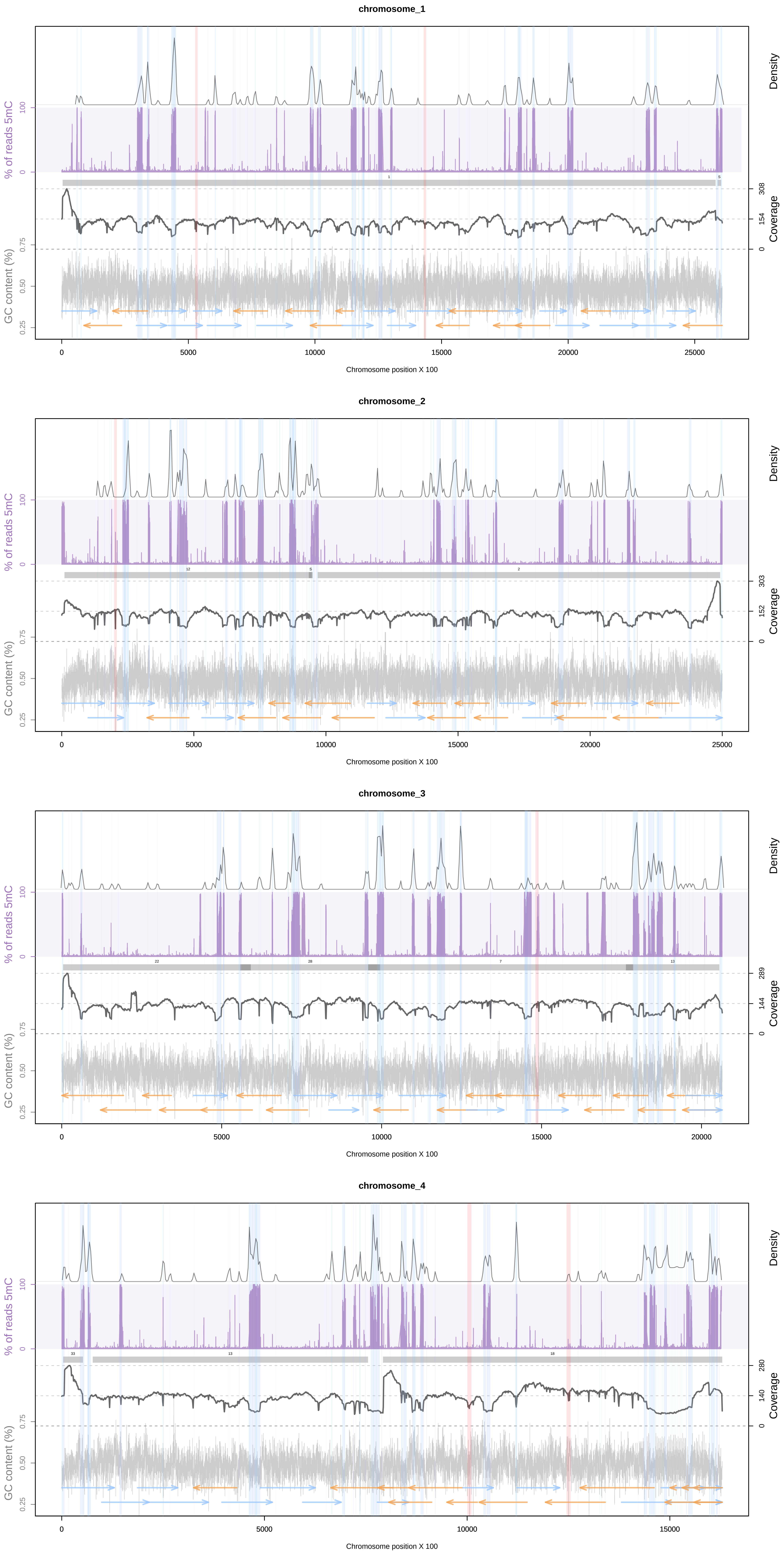

chromosome_1
Density
% of reads 5mC
100
0
Coverage
308
154
0
GC content (%)
0.75
0.50
0.25
0 5000 10000 15000 20000 25000
Chromosome position X 100
chromosome_2
Density
% of reads 5mC
100
0
12
S
2
Coverage
303
152
0
GC content (%)
0.75
0.50
0.25
0 5000 10000 15000 20000 25000
Chromosome position X 100
chromosome_3
Density
% of reads 5mC
100
0
22
28
7
13
Coverage
289
144
0
GC content (%)
0.75
0.50
0.25
0 5000 10000 15000 20000
Chromosome position X 100
chromosome_4
Density
% of reads 5mC
100
0
33
13
18
Coverage
280
140
0
GC content (%)
0.75
0.50
0.25
0 5000 10000 15000
Chromosome position X 100

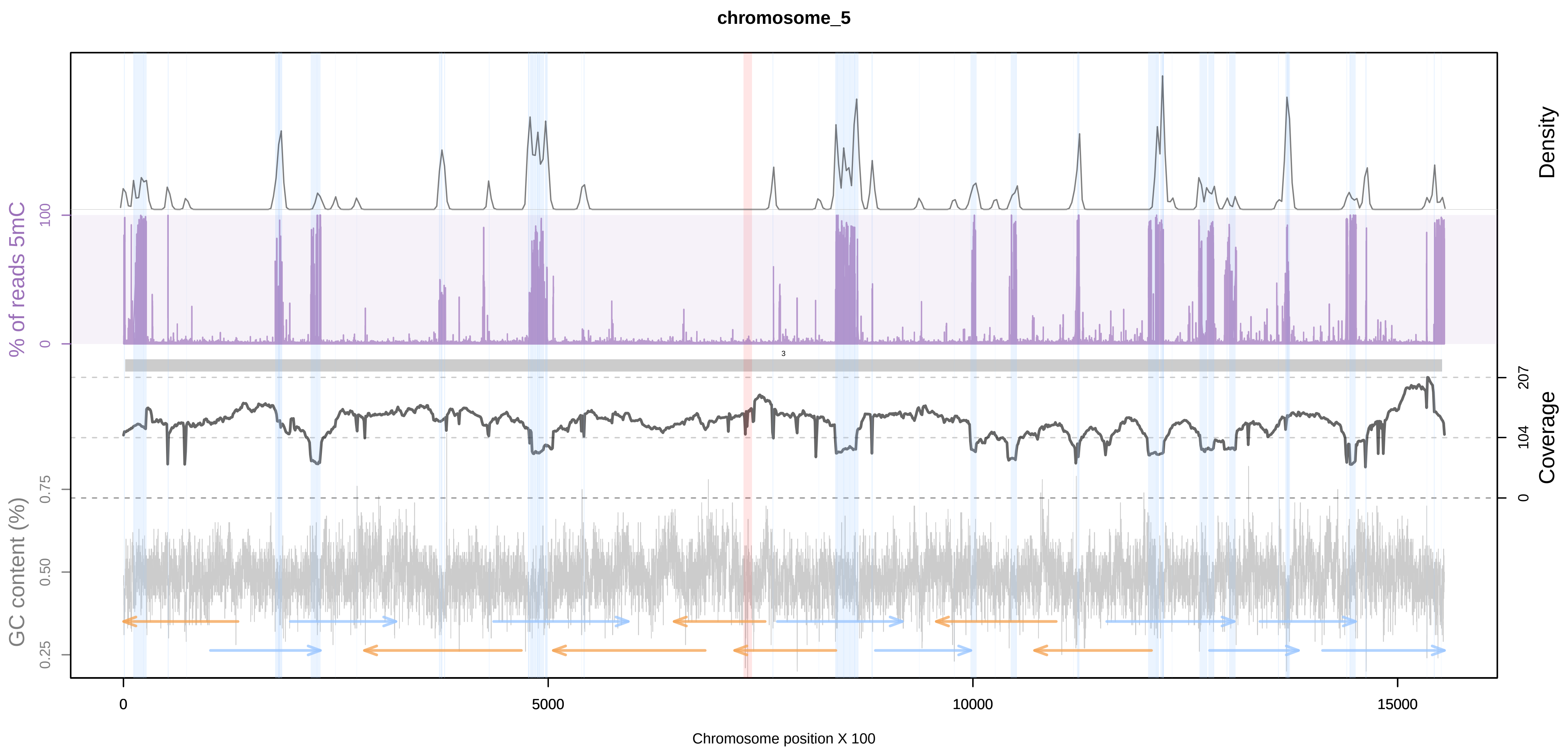
chromosome_5
Density
% of reads 5mC
100
0
3
207
104
0
Coverage
GC content (%)
0.75
0.50
0.25
0
5000
10000
15000
Chromosome position X 100

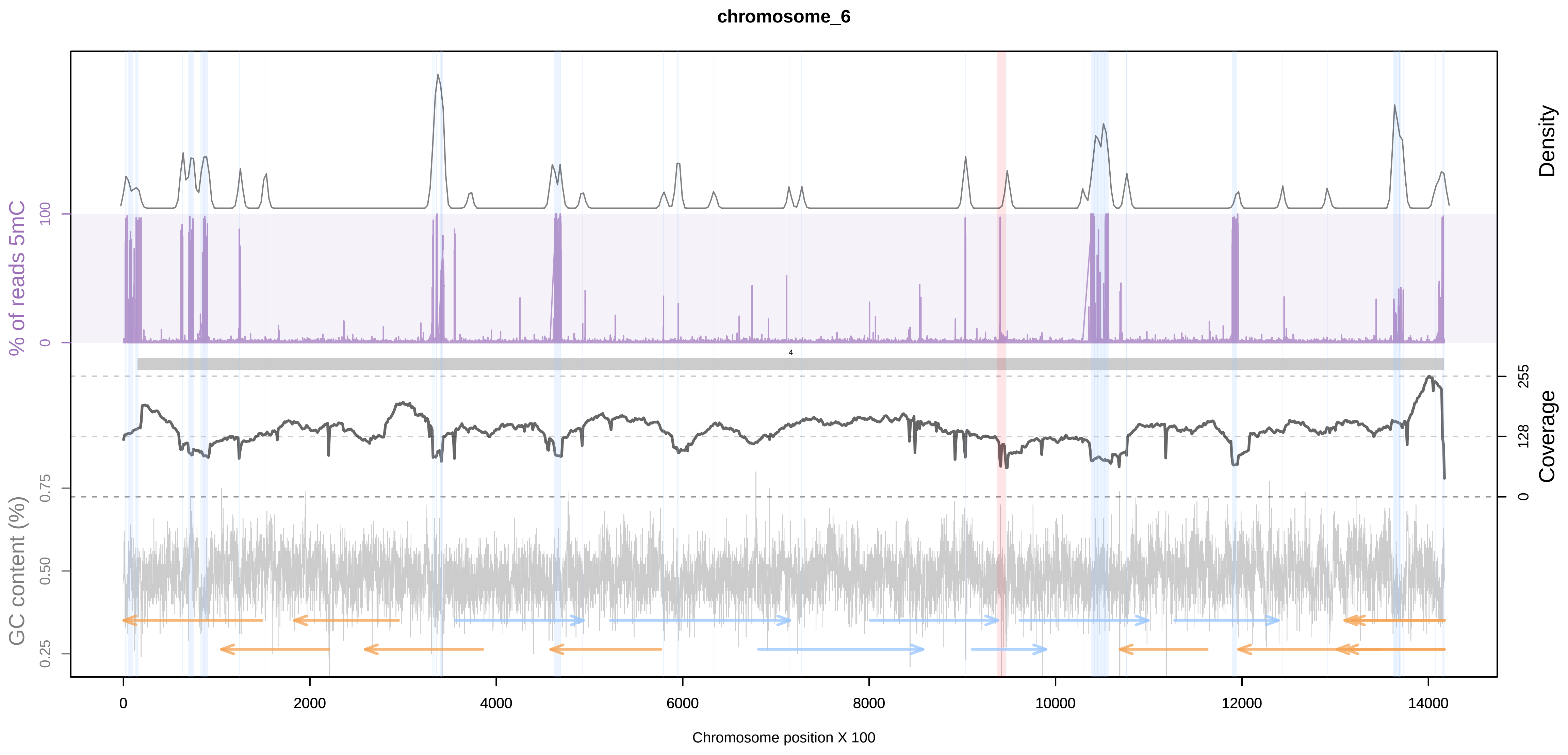
chromosome_6
Density
% of reads 5mC
100
0
4
255
128
0
Coverage
GC content (%)
0.75
0.50
0.25
0
2000
4000
6000
8000
10000
12000
14000
Chromosome position X 100

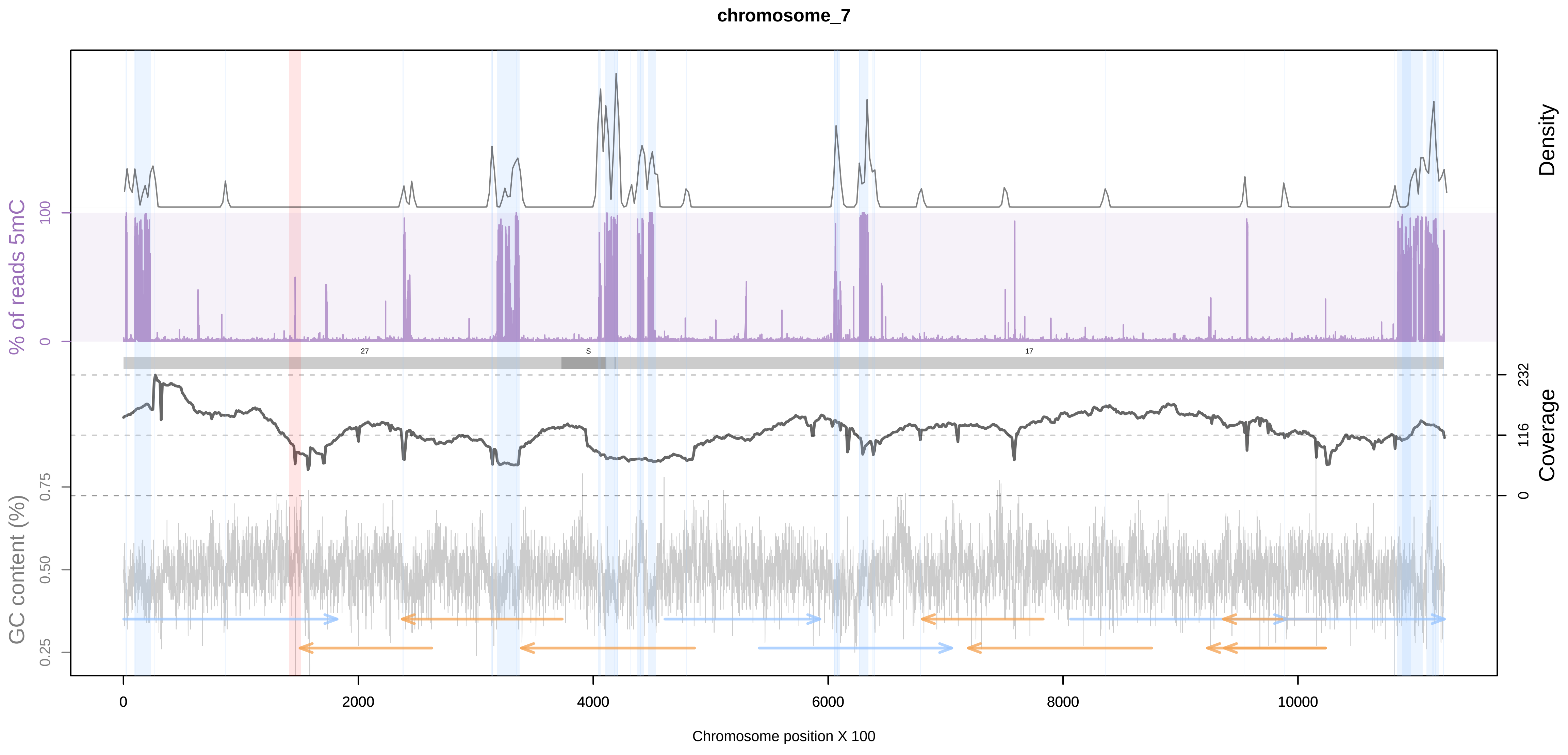
chromosome_7
Density
% of reads 5mC
100
0
27
S
17
232
116
0
Coverage
GC content (%)
0.75
0.50
0.25
0
2000
4000
6000
8000
10000
Chromosome position X 100

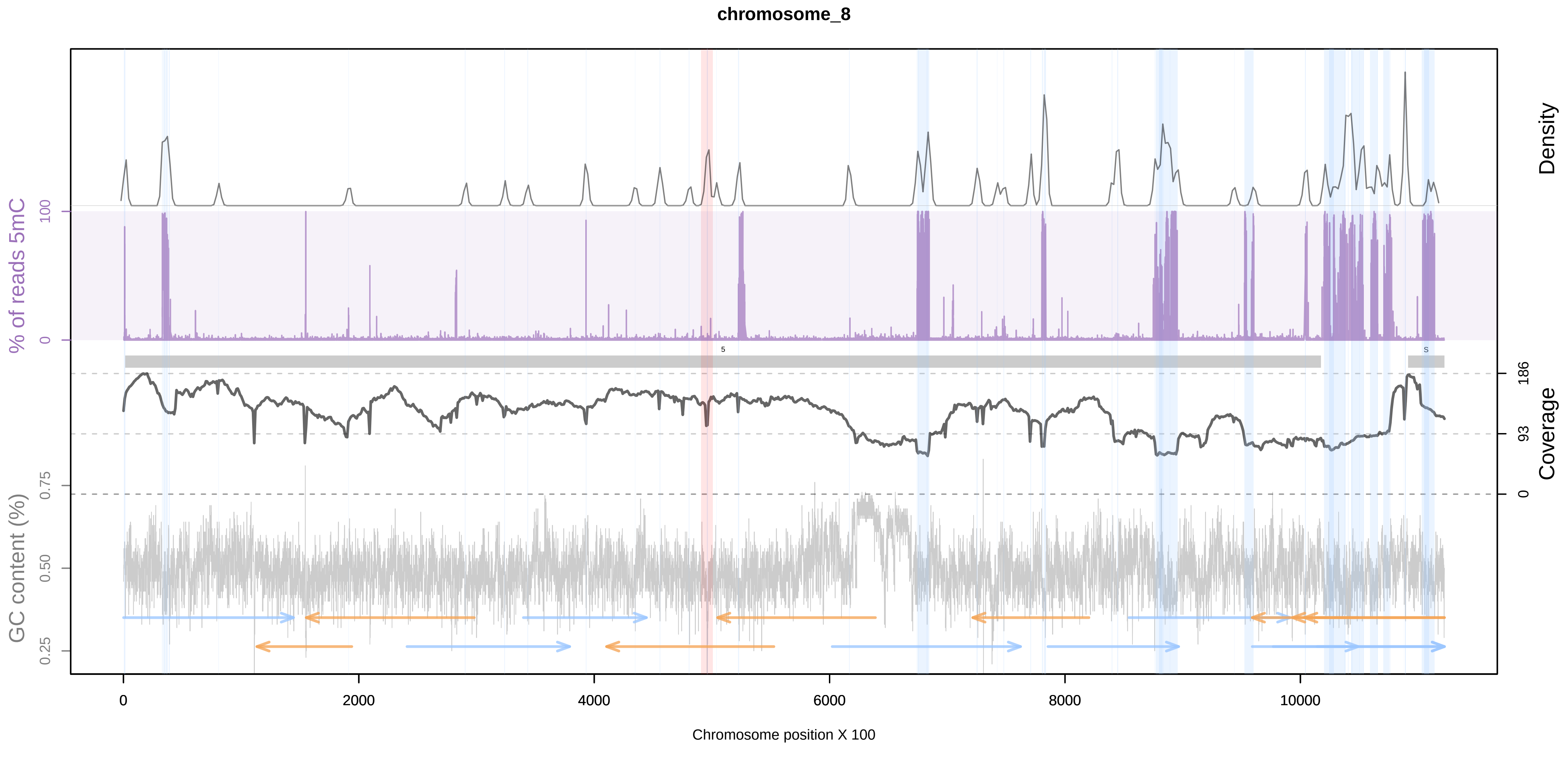
chromosome_8
Density
% of reads 5mC
100
0
5
S
186
93
0
Coverage
GC content (%)
0.75
0.50
0.25
0
2000
4000
6000
8000
10000
Chromosome position X 100

chromosome_9

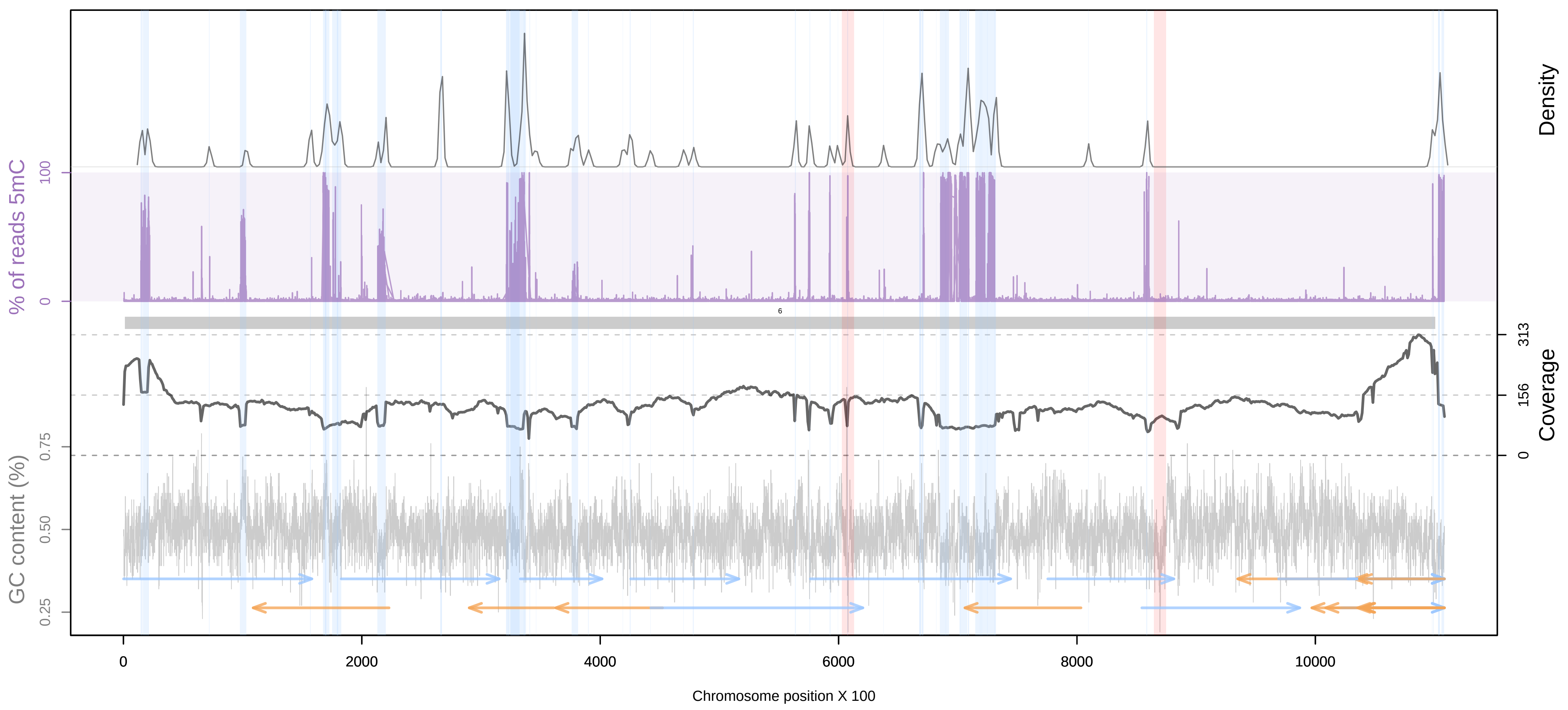


chromosome_10

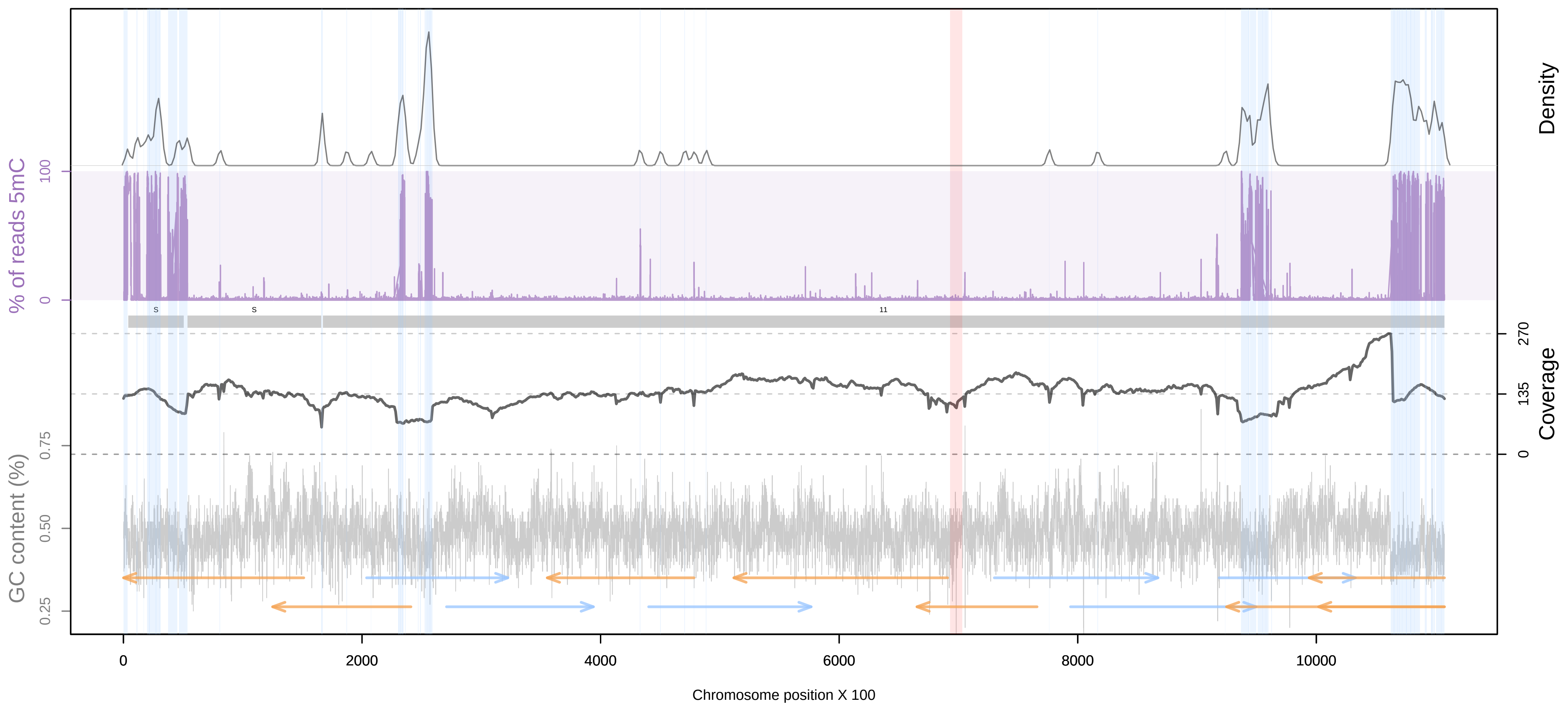


chromosome_11

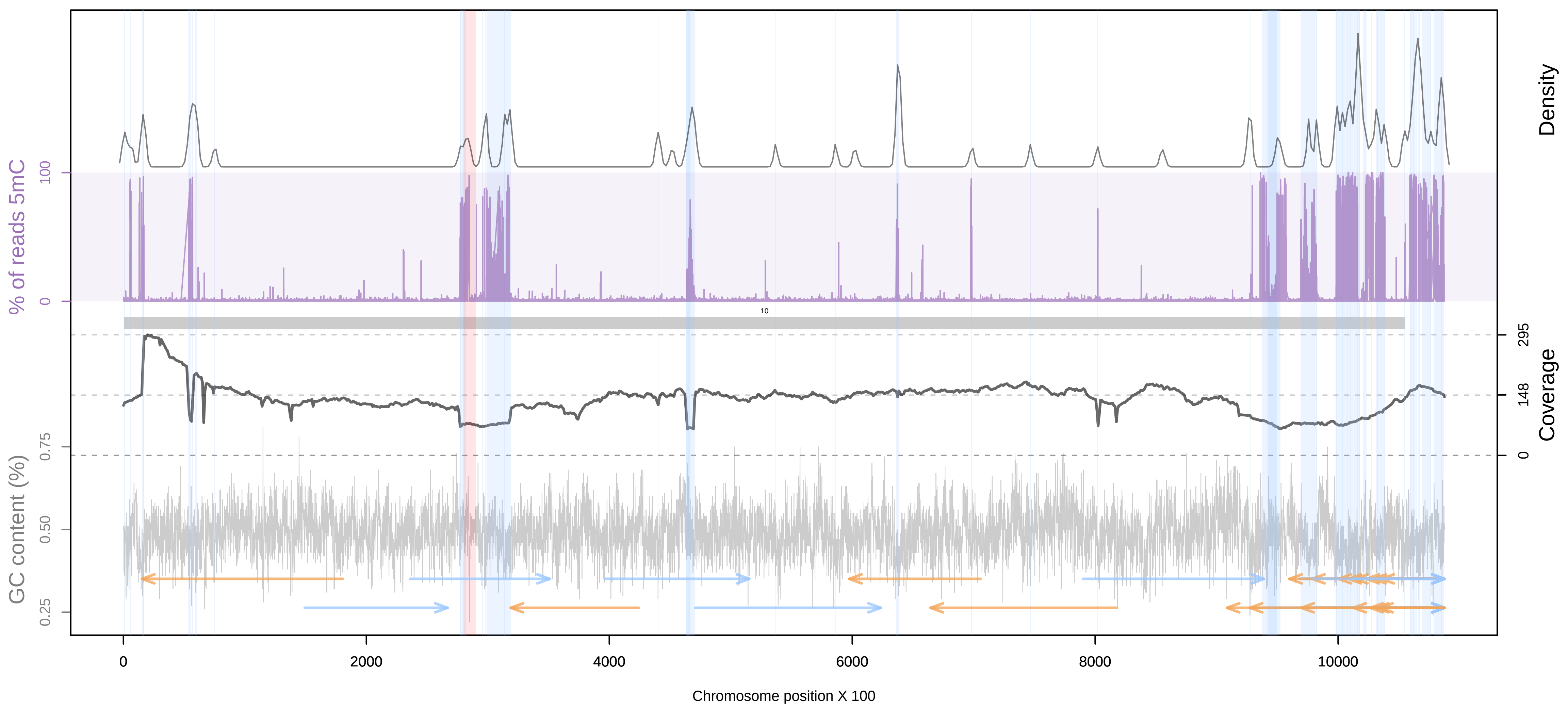


chromosome_12

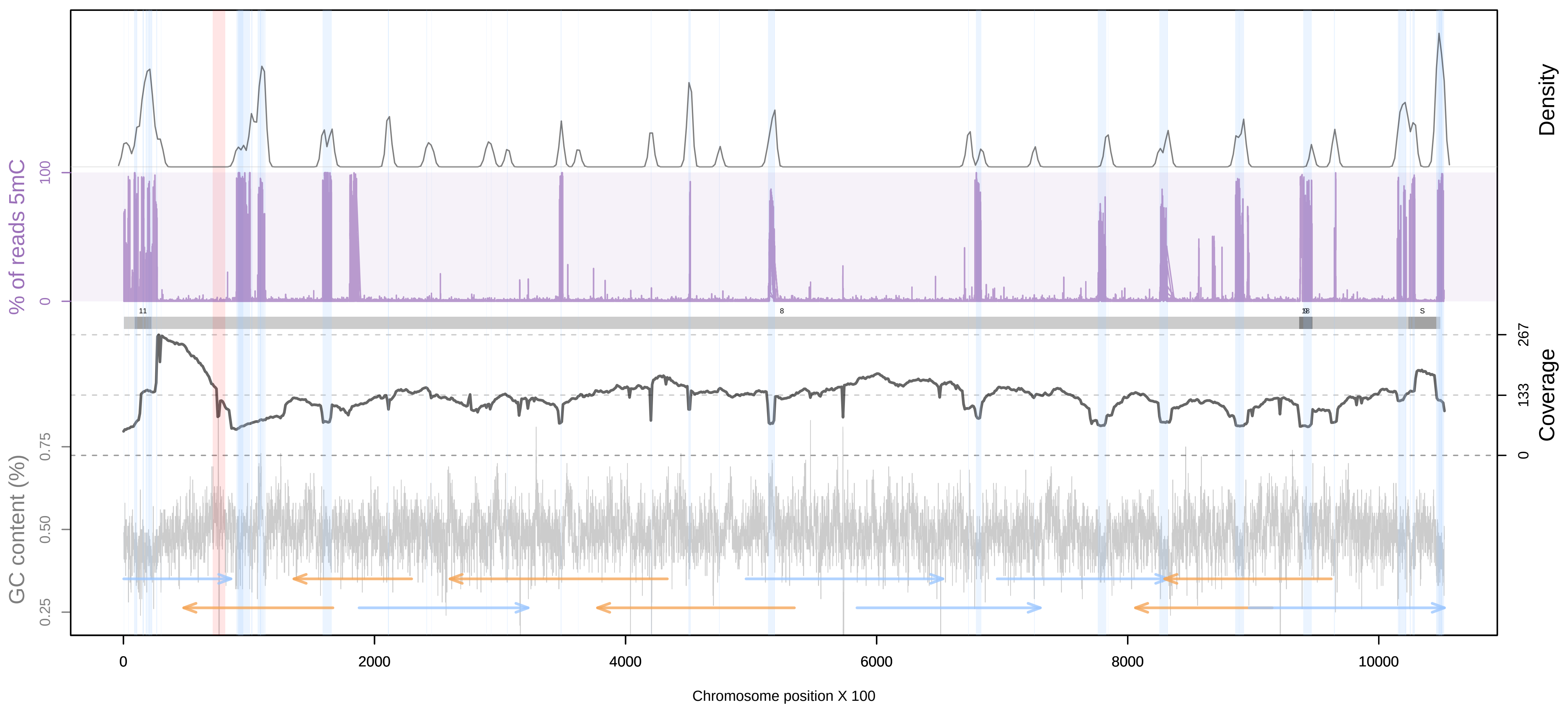

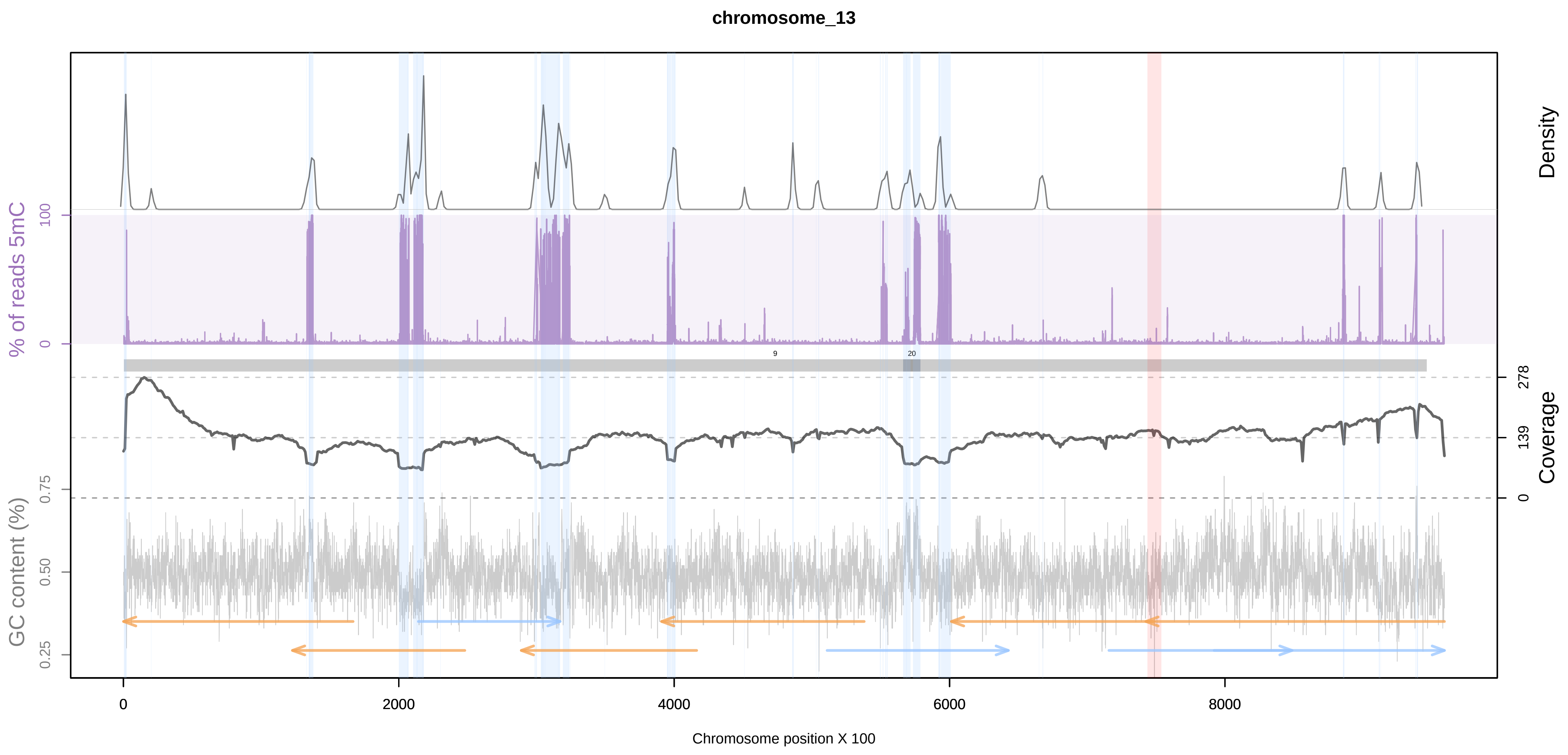
chromosome_13
Density
% of reads 5mC
100
0
278
139
0
Coverage
GC content (%)
0.75
0.50
0.25
0
2000
4000
6000
8000
Chromosome position X 100

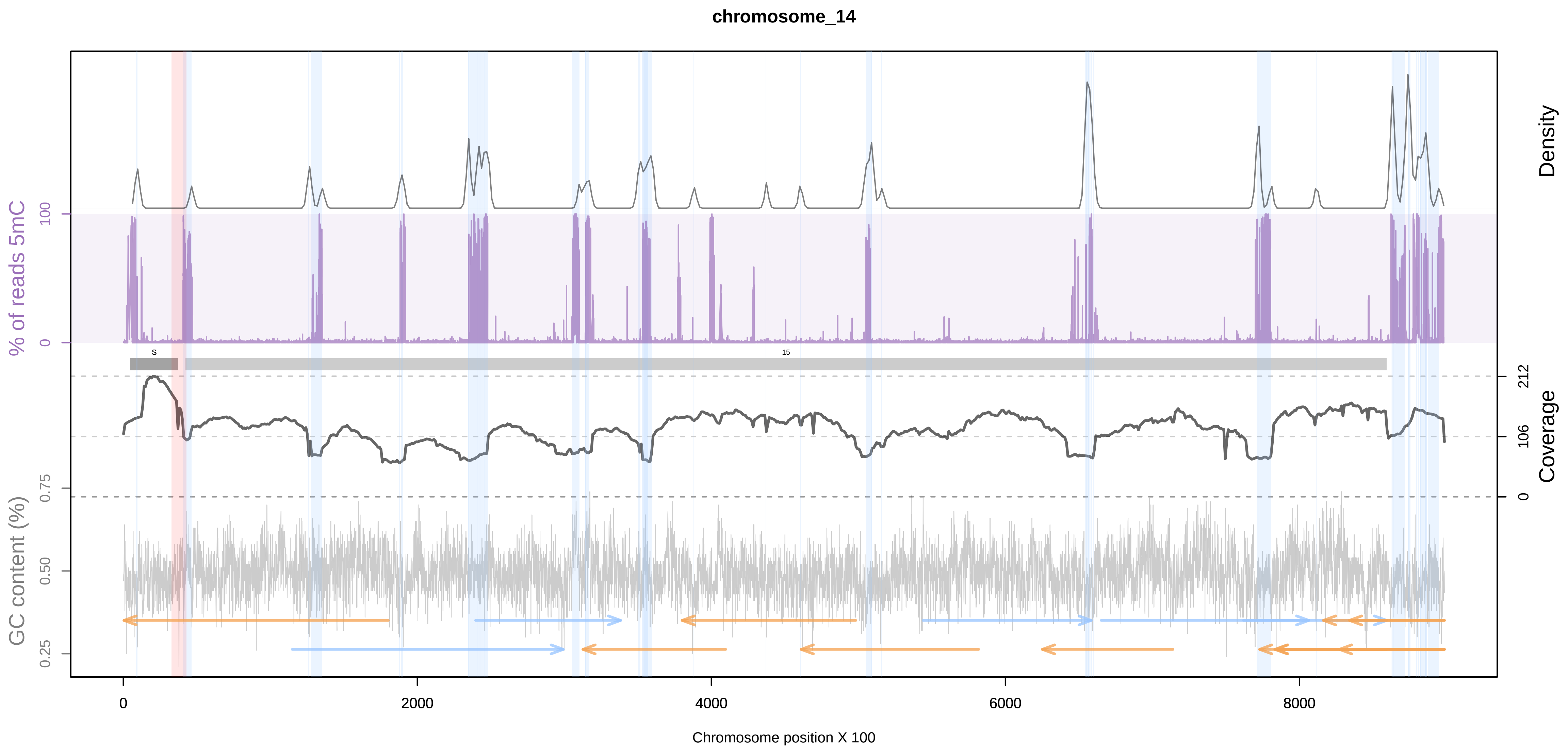
chromosome_14
Density
% of reads 5mC
100
0
212
106
0
Coverage
GC content (%)
0.75
0.50
0.25
0
2000
4000
6000
8000
Chromosome position X 100

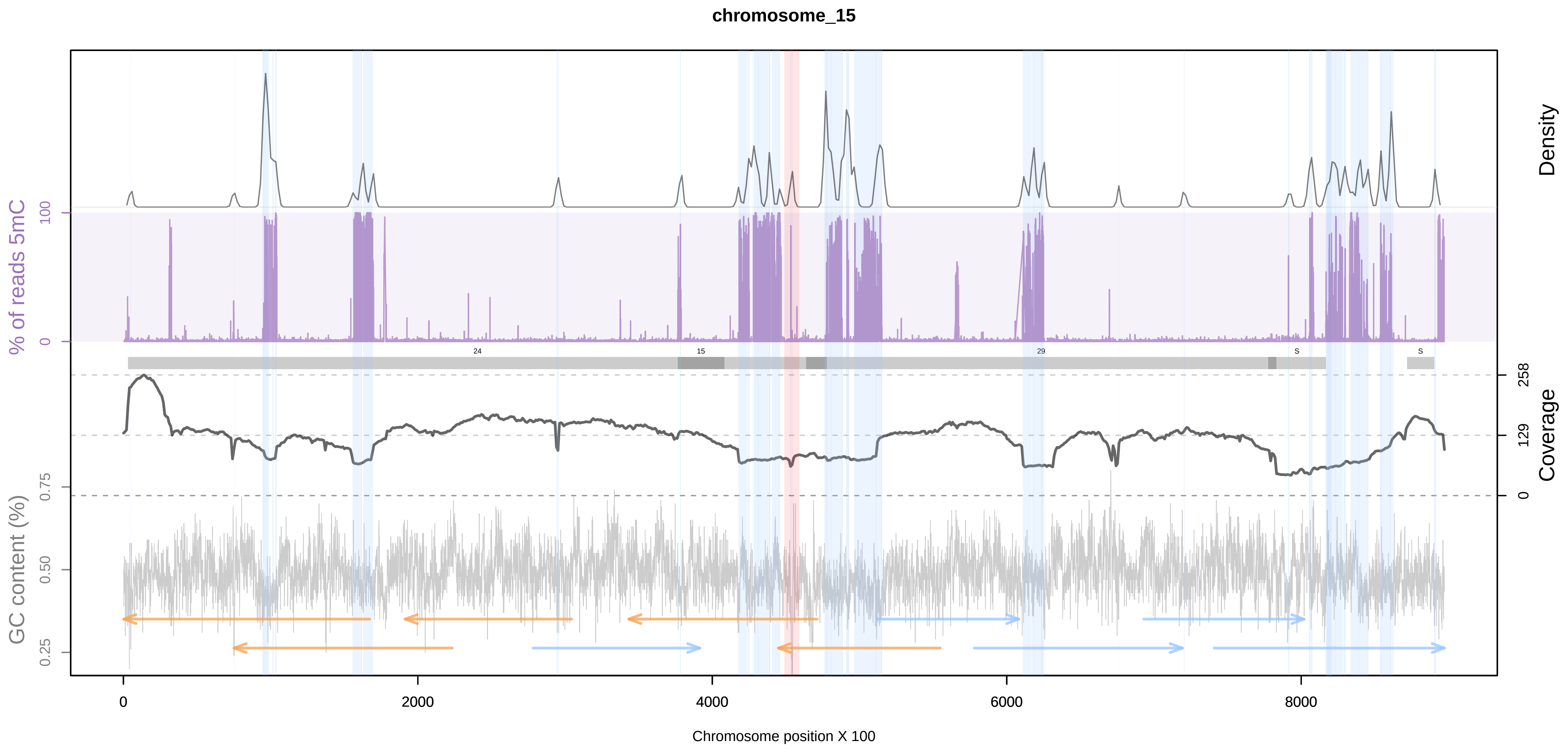
chromosome_15
Density
% of reads 5mC
100
0
258
129
0
Coverage
GC content (%)
0.75
0.50
0.25
0
2000
4000
6000
8000
Chromosome position X 100

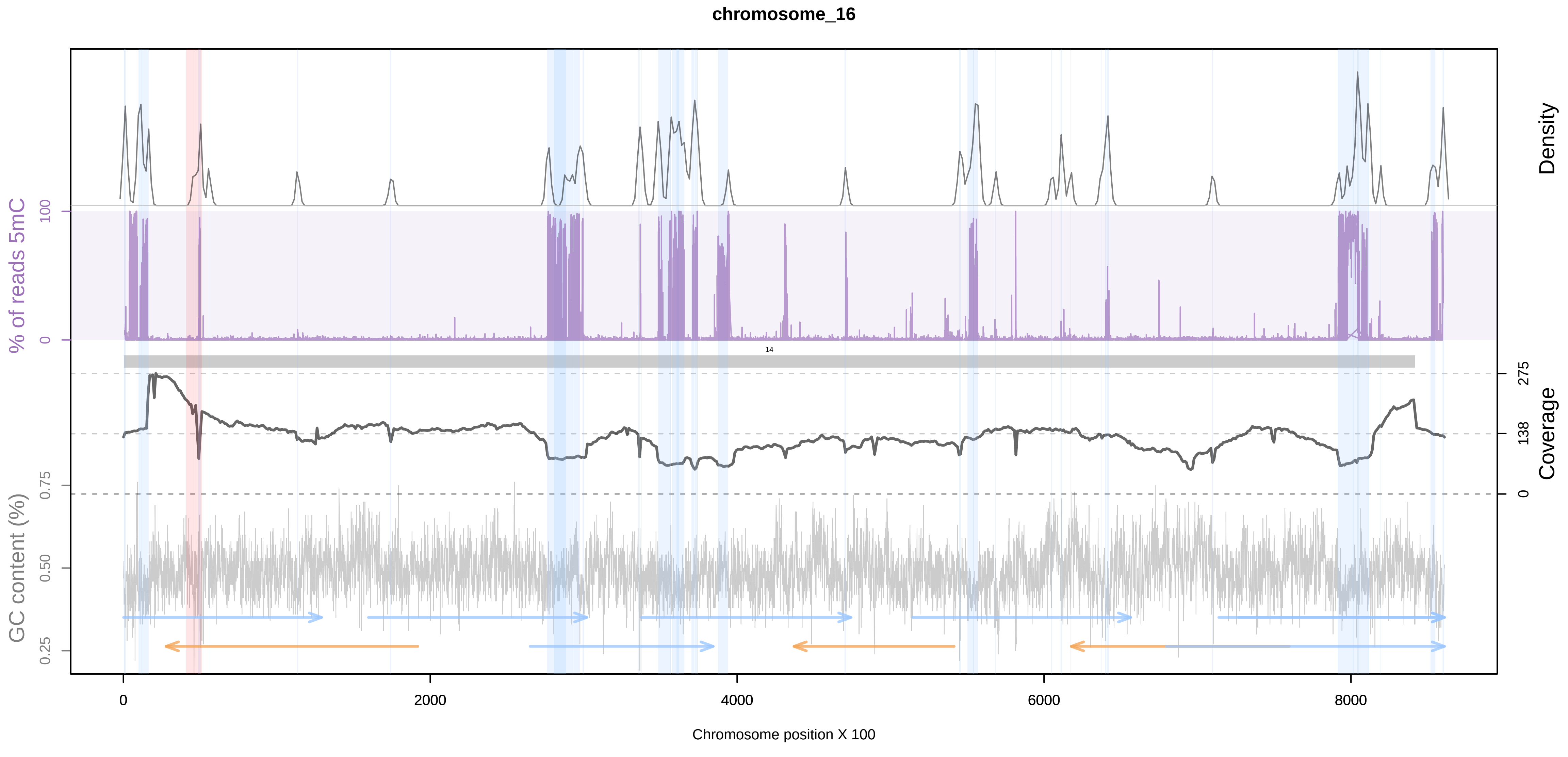
chromosome_16
Density
% of reads 5mC
100
0
275
138
0
Coverage
GC content (%)
0.75
0.50
0.25
0
2000
4000
6000
8000
Chromosome position X 100

chromosome_17

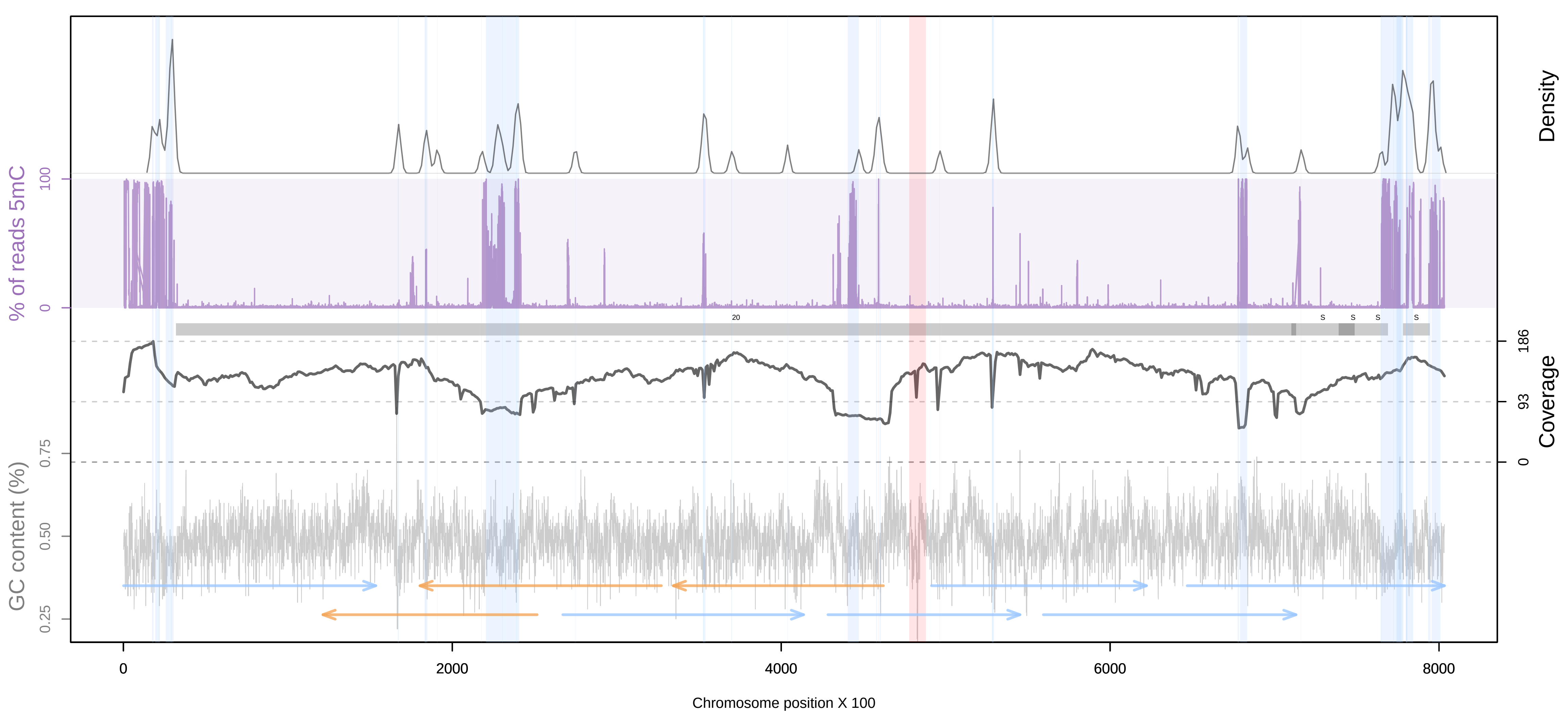


chromosome_18

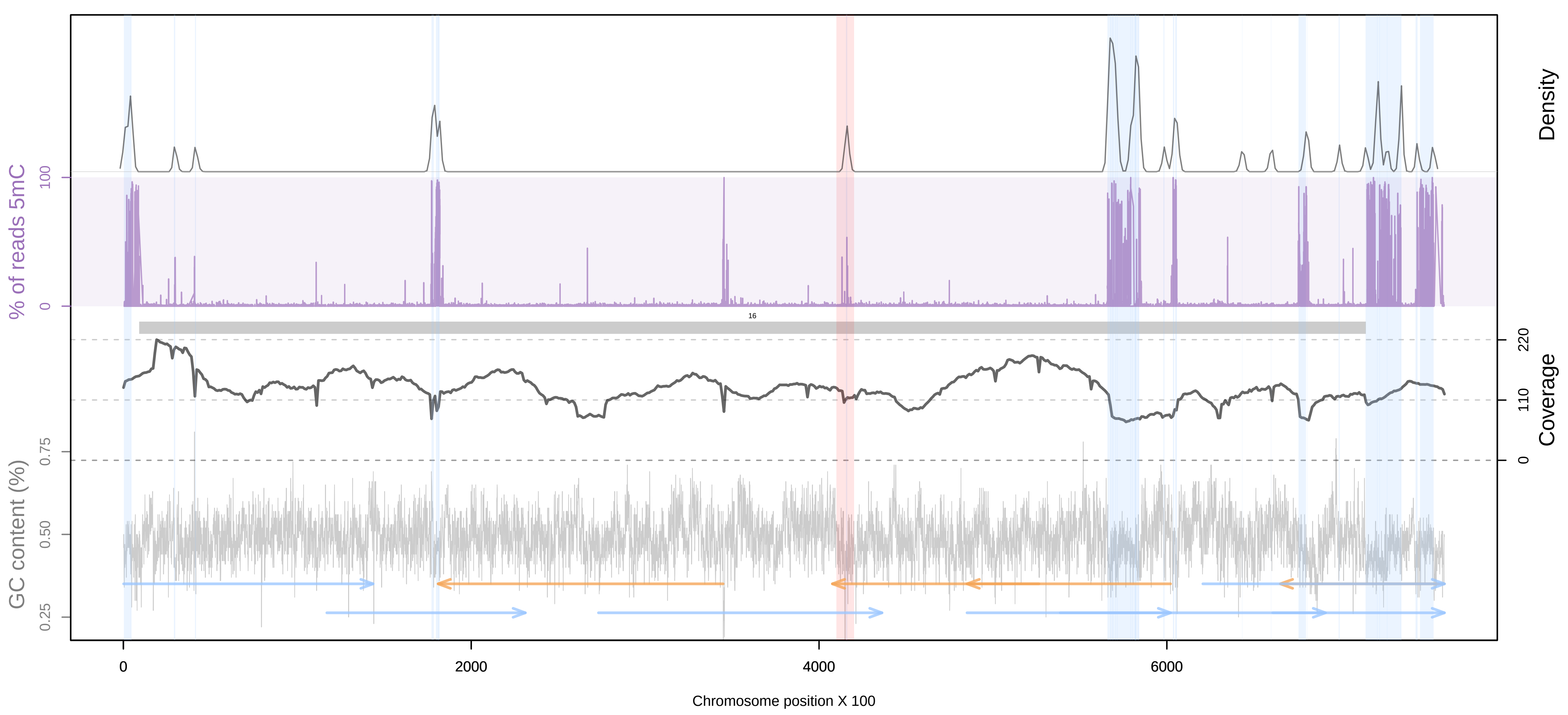


chromosome_19

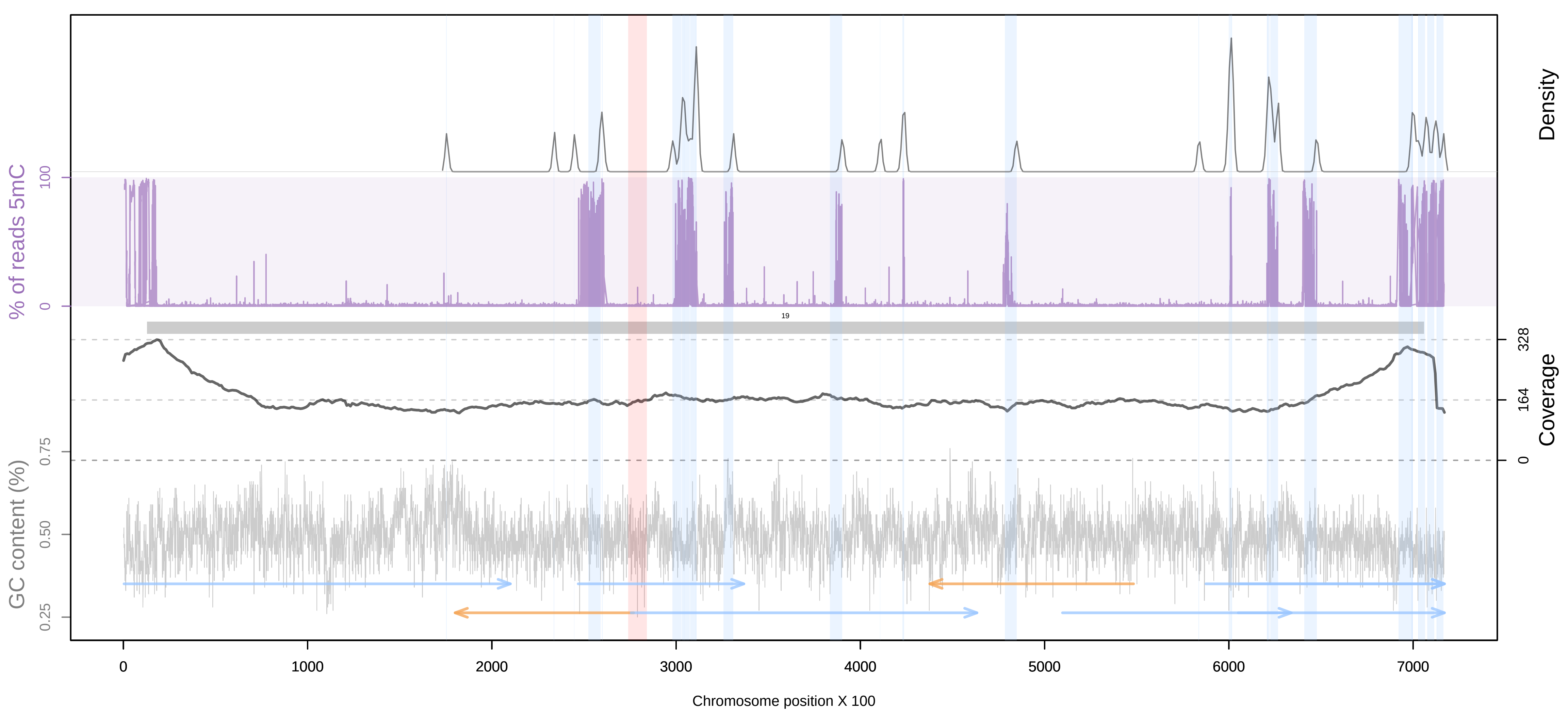


chromosome_20

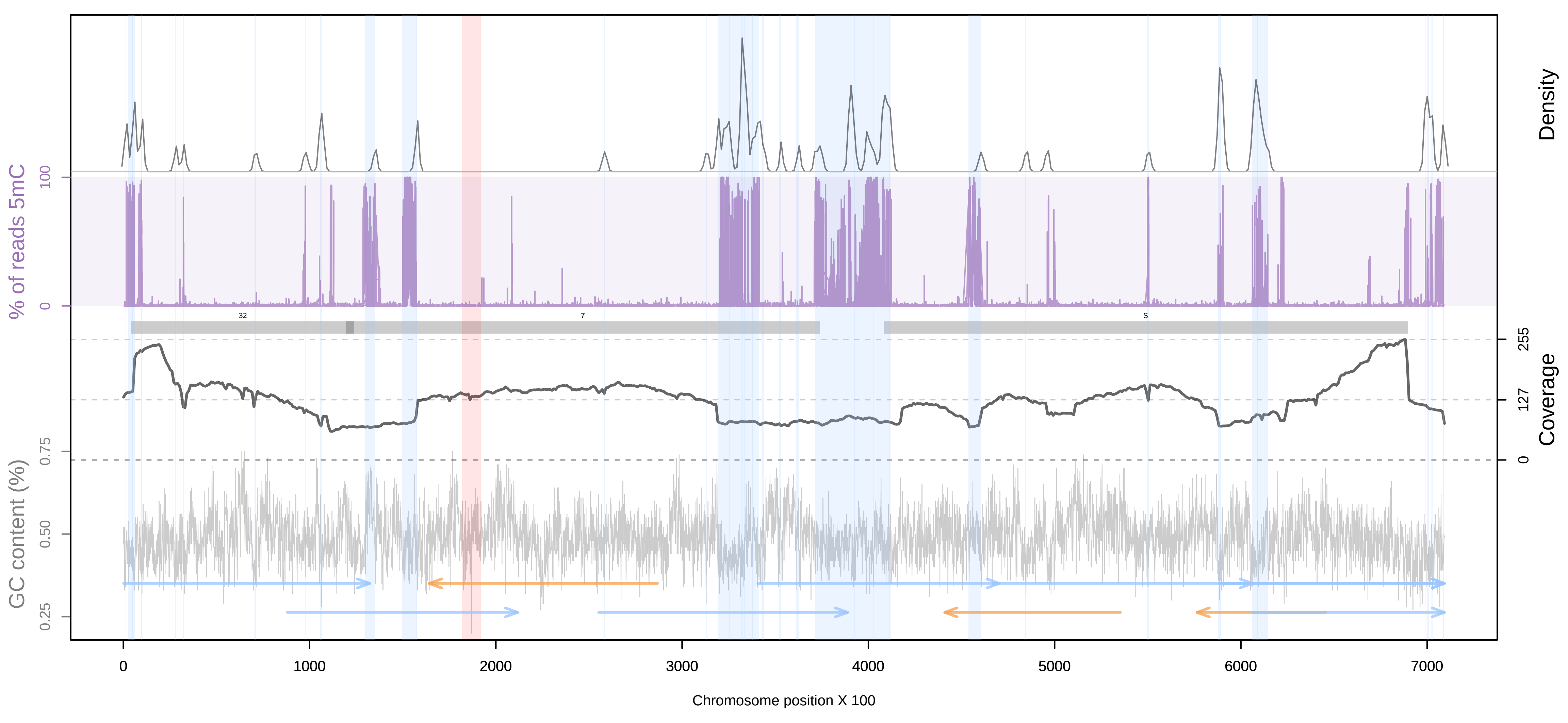

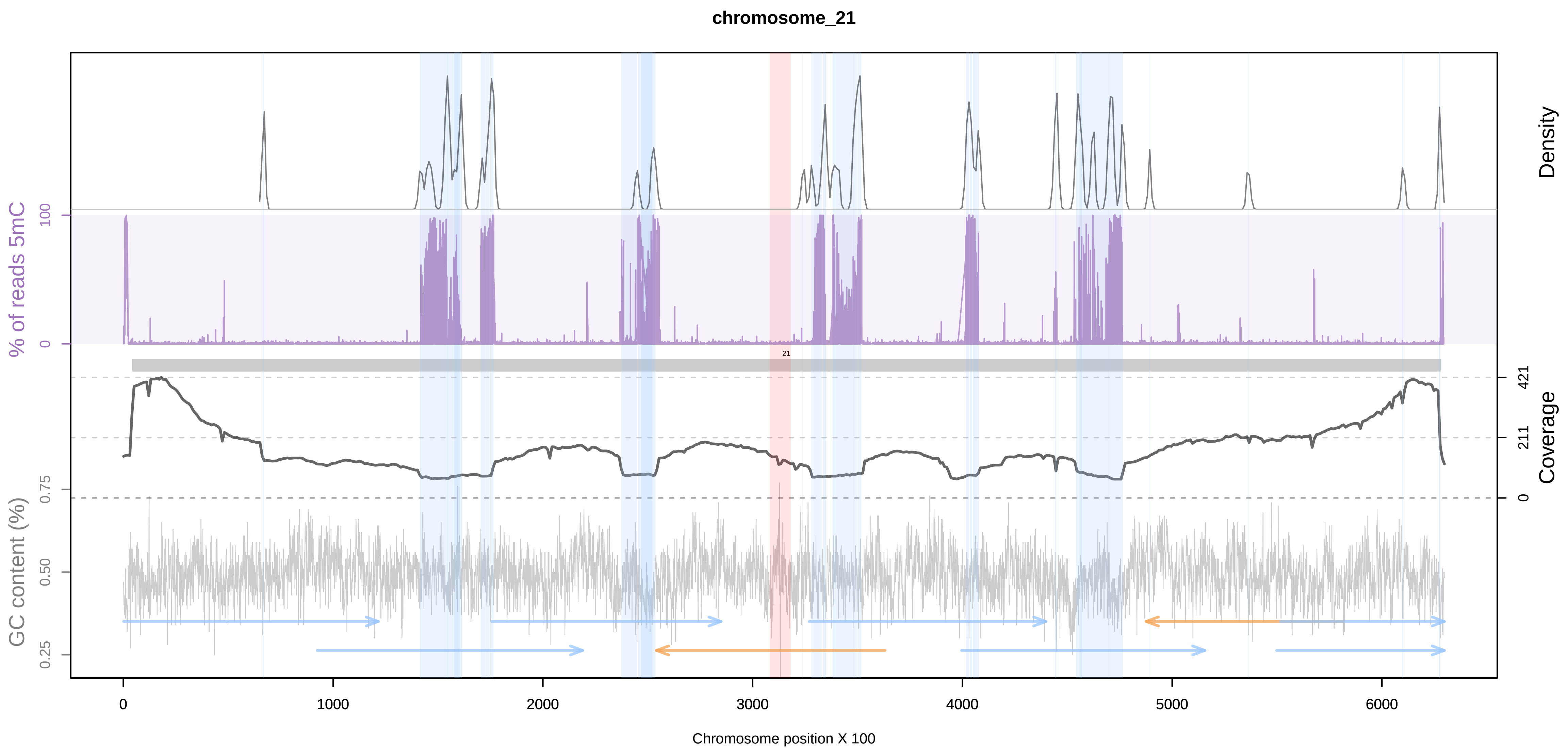
chromosome_21
Density
% of reads 5mC
100
0
Coverage
421
211
0
GC content (%)
0.75
0.50
0.25
0
1000
2000
3000
4000
5000
6000
Chromosome position X 100

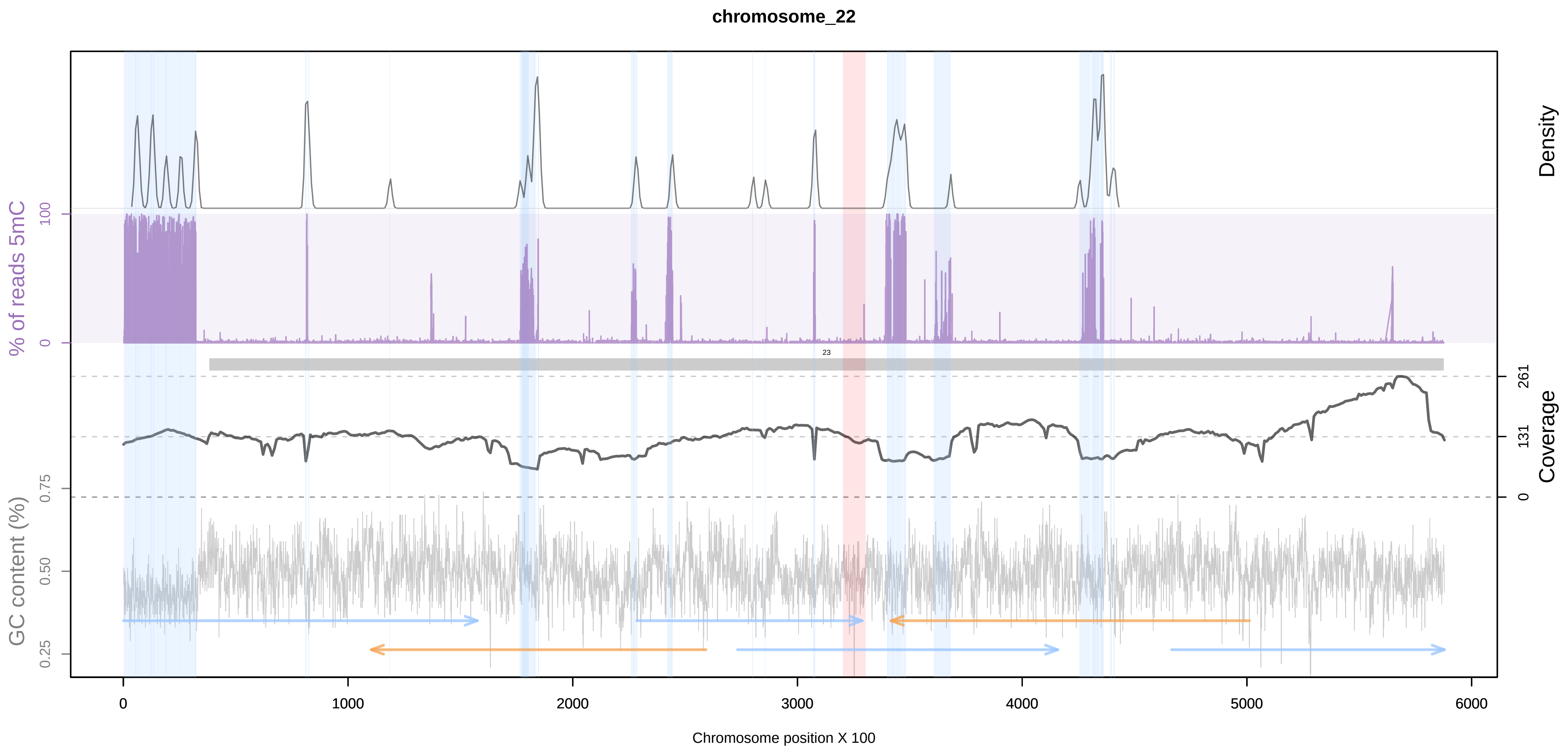
chromosome_22
Density
% of reads 5mC
100
0
Coverage
261
131
0
GC content (%)
0.75
0.50
0.25
0
1000
2000
3000
4000
5000
6000
Chromosome position X 100

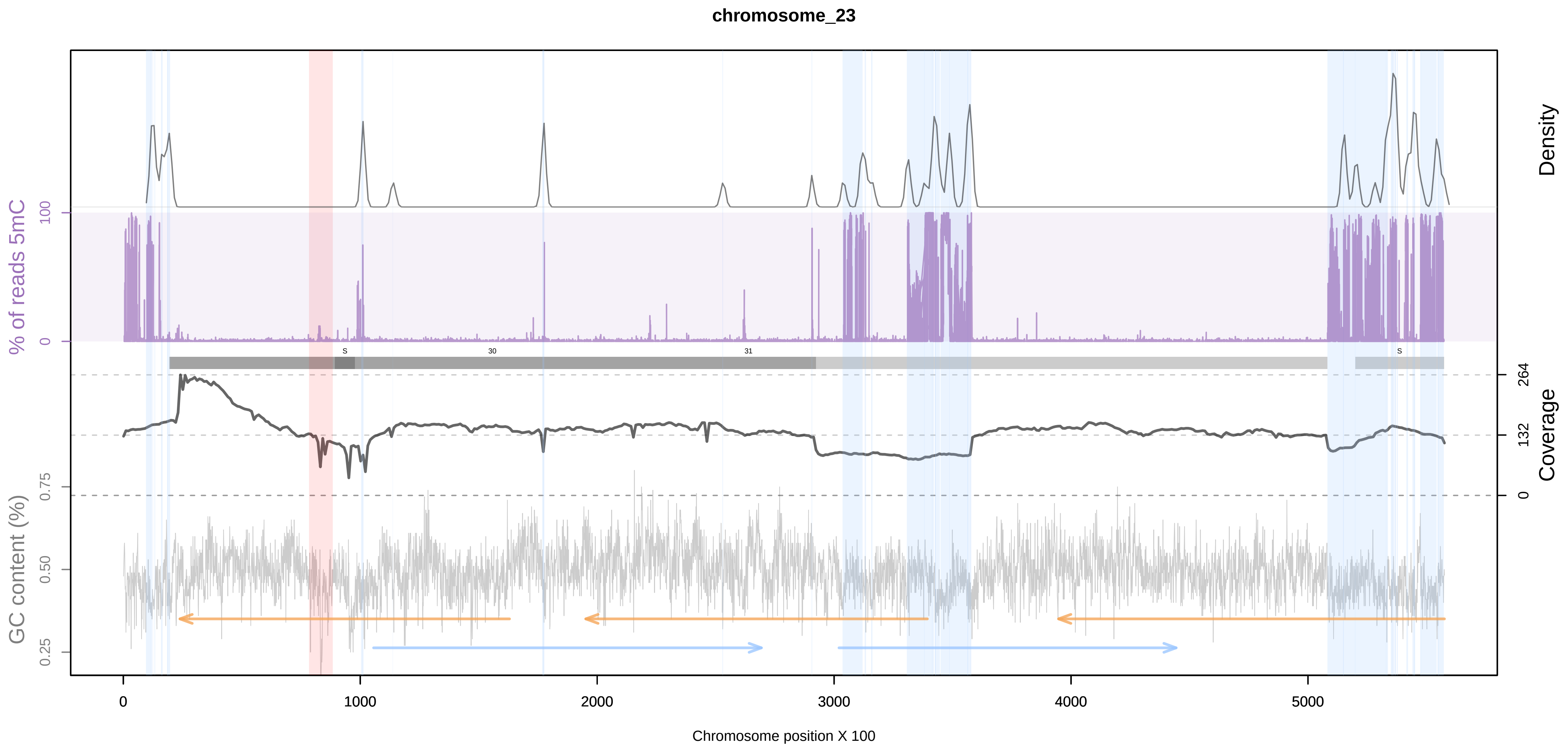
chromosome_23
Density
% of reads 5mC
100
0
Coverage
264
132
0
GC content (%)
0.75
0.50
0.25
0
1000
2000
3000
4000
5000
Chromosome position X 100

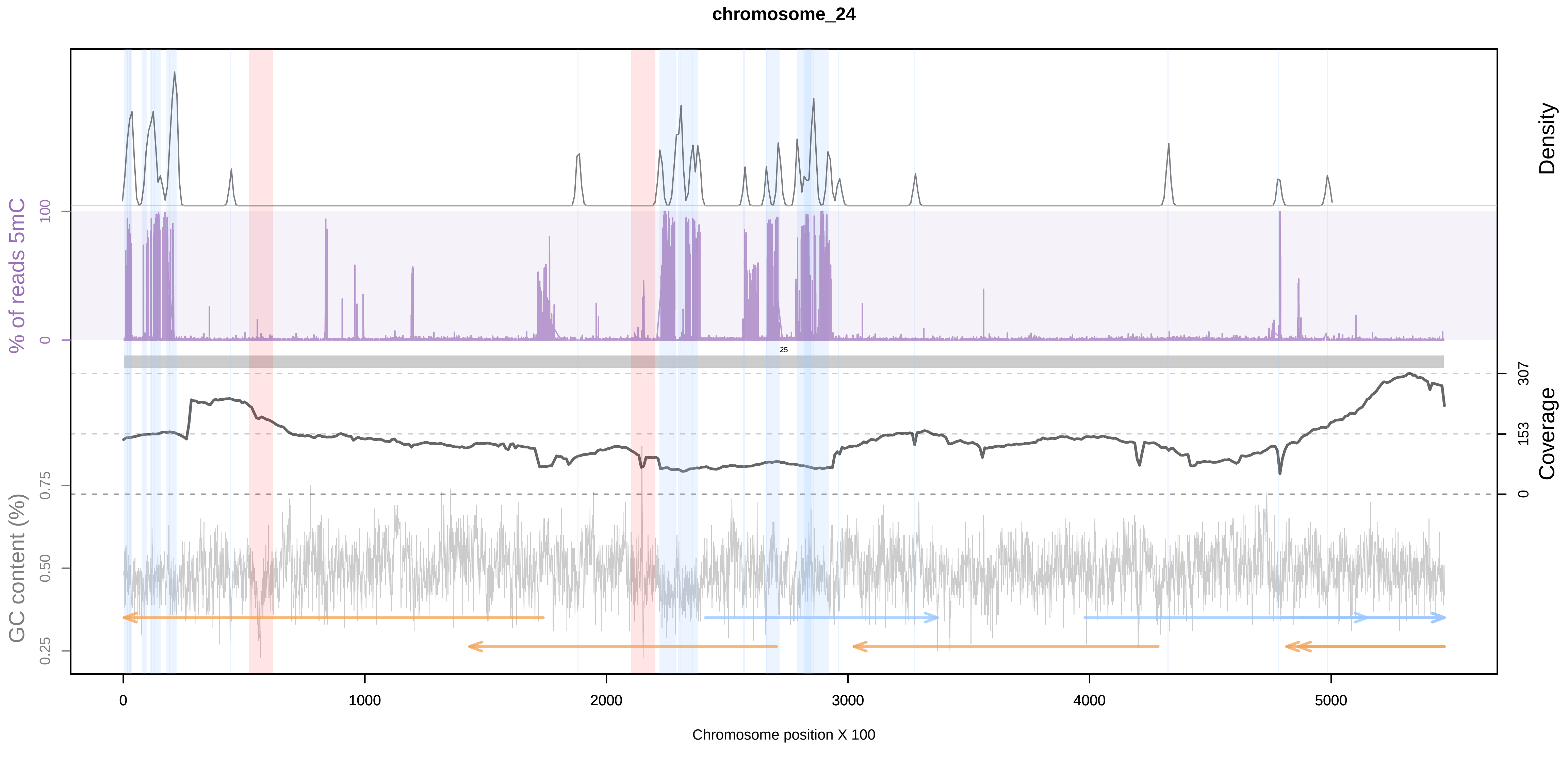
chromosome_24
Density
% of reads 5mC
100
0
Coverage
307
153
0
GC content (%)
0.75
0.50
0.25
0
1000
2000
3000
4000
5000
Chromosome position X 100

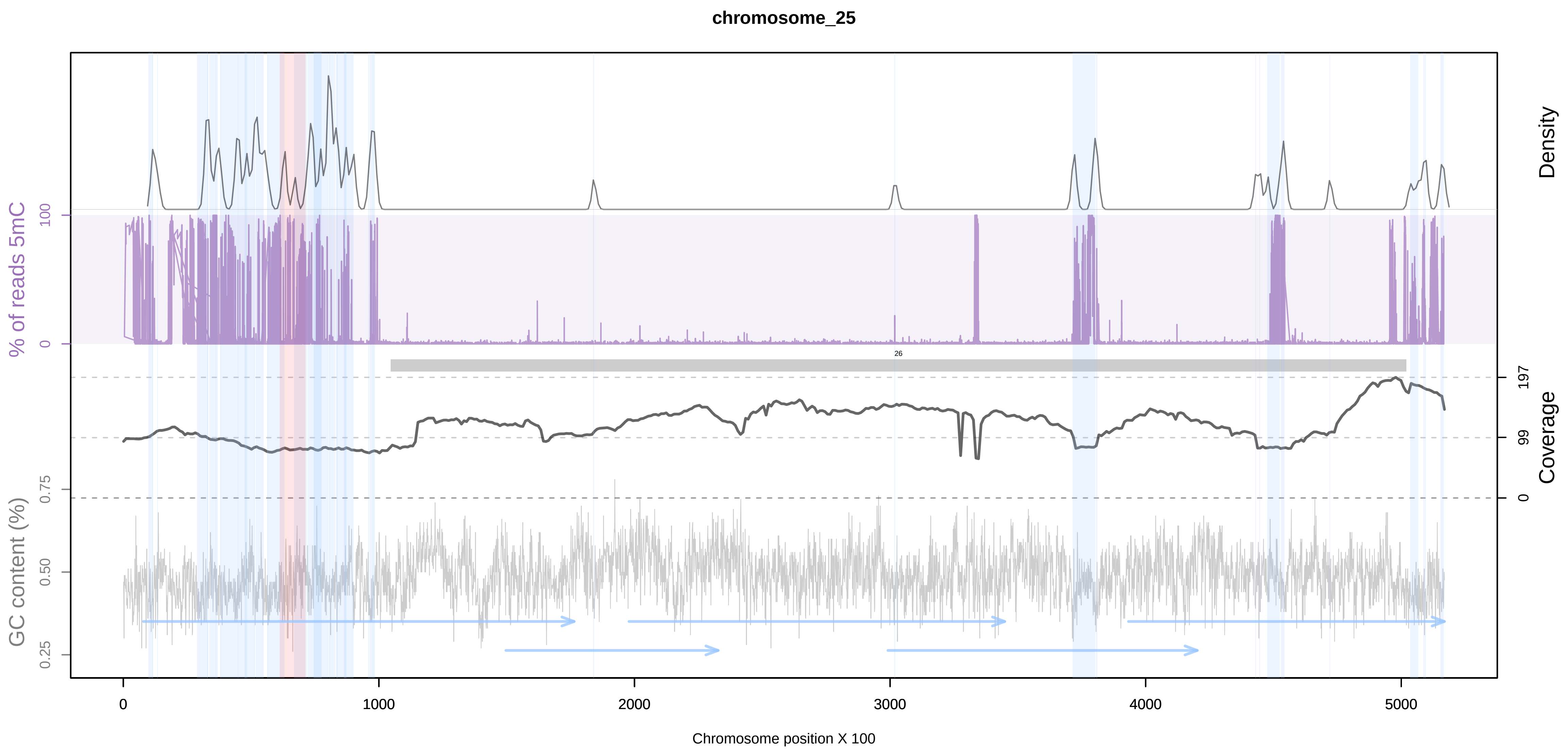
chromosome_25
Density
% of reads 5mC
100
0
26
197
99
0
Coverage
GC content (%)
0.75
0.50
0.25
0
1000
2000
3000
4000
5000
Chromosome position X 100
