## Supplementary figures and images for "Telomere-to-telomere genome assembly of *Phaeodactylum tricornutum*"

### Supplemental Figure 1

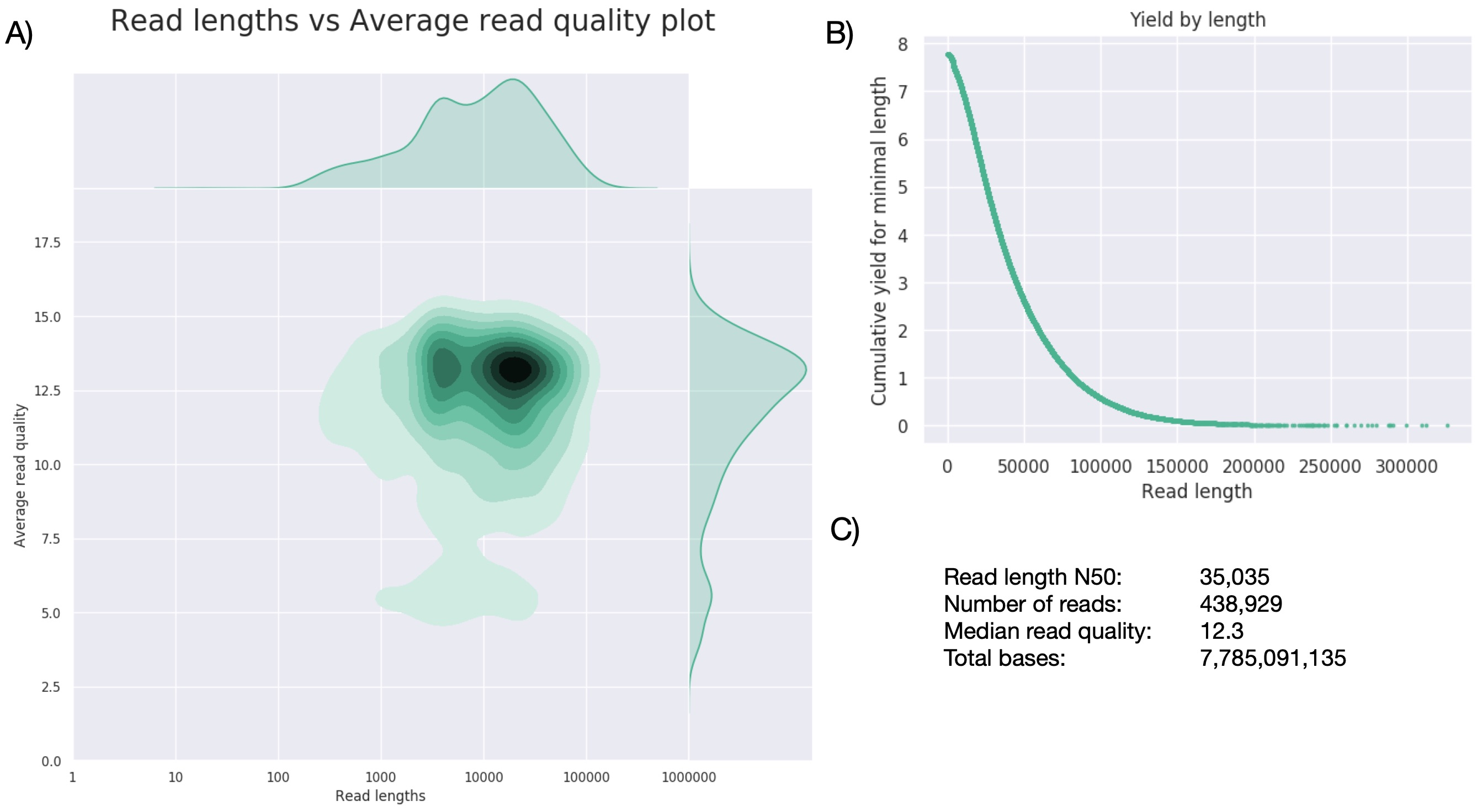
