## Supplemental methods 6 for "Telomere-to-telomere genome assembly of *Phaeodactylum tricornutum*"

__A summary of the steps used to polish the pt genome__

A pipeline was used for initial polishing of the raw, wild type, haplotype *Phaeodactylum tricornutum* genome assembly. This pipelines was based on a previously described pipeline used to polish the genomes of metagenomically assembled prokaryotes [@Giguere:2020bn]. The pipline used to polish *P. tricornutum* consisted of four rounds of Rebaler (Ryan Wick) followed by two rounds of medaka-consensus (Oxford Nanopore) and finally two rounds of Pilon [@Walker:2014ky] in hybrid mode.

Rebaler was used to polish the raw assembly directly with all long reads using default parameters. The polished from each Rebaler run was used as the input for subsequent Rebaler runs until the genome had been polished four times. After the fourth round of Rebaler polishing, the raw genome was polished by medaka-consensus with default parameters using all long reads, twice. 

Long and short reads were mapped to the genome that was polished from medaka-consensus using minimap2 [@Li:2016fb] and bowtie2 [@Langmead:2012jh], respectively. Long reads were mapped using minimap2 with parameters ```-aLQx map-ont --secondary=no --sam-hit-only```. The mapped reads were then filtered using Gerenuq (github.com/abahcheli/gerenuq) with default parameters. Short, Illumina reads were mapped using bowtie2 with parameters ```--no-unal --no-discordant --end-to-end --very-sensitive```. Mapped Illumina reads were filtered, sorted and indexed using samtools view [@Li:2009ka] with parameters ```-F 256```. Long and short mapped reads were used in Pilon to polished the medaka-polished genome with parameters ```-Xmx32G -jar --fix snps,indels```. This mapping, filtering and polishing with pilon process was repeated twice for the genome. 

Polished genome quality was interrogated by analyzing coverage maps. Long reads were mapped to both the unpolished and the polished genomes using minimap2, and mapped reads were filtered with Gerenuq, as described above. Filtered long read coverage was used to determine which chromosomes were supported by consistent long-read coverage and which chromosomes required further polishing. Unfortunately, after a single round of polishing, most polished chromosomes required further polishing due to regions of insufficient coverage and many polished chromosomes actually showed worse coverage plots than the unpolished versions. 

One major issue that continues to impair long-read polishing and assembly strategies is the mismapping of long-reads to repetitive or polymorphic loci. This is especially a problem during the haploid assembly of polyploid genomes, where loci are polymorphic from both haplotype and population divergence. One solution to this problem is described previously [@Giguere:2020Bn] and involves filtering mapped reads by a number of parameters to improve mapping quality and increase confidence in read alignments. 

We implemented Gerenuq into the medaka-consensus pipeline to filter mapped reads prior to medaka's polishing and neural network consensus steps. The modified medaka-consensus pipeline was used to polish exclusively the chromosomes whose coverage maps indicated poor assembly quality. Any chromosomes with drops in coverage to zero were manually inspected and corrected using long reads that span the region.
