## Supplemental code 7 for "Telomere-to-telomere genome assembly of *Phaeodactylum tricornutum*"

Supplemental code for Telomere-to-telomere genome assembly of *Phaeodactylum tricornutum*

Extracting telomere-containing ultra long reads:

```
gunzip -c pt_guppy-3-6.fastq.gz | NanoFilt -l 50000 \ 
    grep -A 2 -B 1 --no-group-separator \
    -E "AACCCTAACCCTAACCCT|AGGGTTAGGGTTAGGGTT" - \ 
    | gzip > pt_telomeric_reads.fastq.gz
```

All-vs-all mapping of telomere containing reads: 

```
minimap2 -x ava-ont -t 40 pt_telomeric_reads.fastq.gz \
    pt_telomeric_reads.fastq.gz > telomere_overlaps.paf 

R

# filtered the overlaps in R
d <- read.table("telomere_overlaps.paf")

# retain only reads with 95% query coverage
filtered <- d[which( (d$V4 - d$V3) / d$V2 > 0.95),] 

# write to output
write.table(filtered, "telomere_overlaps_filtered.paf", sep = "\t", \
    quote = FALSE, col.names=FALSE, row.names=FALSE)
```

Network graph of telomere all-vs-all mapping:

```
R
library(igraph)

d <- read.table("telomere_overlaps_filtered.paf")

subset <- data.frame(from=d$V1, to=d$V6)
g <- graph_from_data_frame(subset)

clu <- components(g)

final <- vector()

for (i in seq(length(groups(clu)))) {
    
    final[i] <- length(groups(clu)[[i]])
    
}

# visualize the coverage per telomere
hist(final, breaks = 50, ylim = c(0, 15), xlim = c(0, 100))
```
